# Limitations of acyclovir and identification of potent HSV antivirals using 3D bioprinted human skin equivalents

**DOI:** 10.1101/2024.12.04.626896

**Authors:** S. Tori Ellison, Ian Hayman, Kristy Derr, Paige Derr, Shayne Frebert, Zina Itkin, Min Shen, Anthony Jones, Wendy Olson, Lawrence Corey, Anna Wald, Christine Johnston, Youyi Fong, Marc Ferrer, Jia Zhu

**Author notes:** These authors contributed equally to this work (S. Tori Ellison, Ian Hayman).

## Abstract

Herpes simplex virus (HSV) infection has worldwide public health concerns and lifelong medical impacts. The standard therapy, acyclovir, has limited efficacy in preventing HSV subclinical virus shedding, and drug resistance occurs in immunocompromised patients, highlighting the need for novel therapeutics. HSV manifests in the skin and mucosal epithelium. Here, we found acyclovir significantly less effective in skin-derived keratinocytes than donor-matched fibroblasts. To recapitulate *in vivo* tissue architecture, we 3D bioprinted human skin equivalents (HSE) in a 96-well plate format amenable for antiviral screening and preclinical testing. We screened a library of 738 compounds with broad targets and mechanisms of action and identified potent antivirals, including 23 known or experimental HSV treatments, validating the translational relevance of our assay. Unlike acyclovir, antivirals against HSV helicase/primase or host replication pathways displayed similar potency across cell types and donor sources in 2D and 3D models. Our 3D bioprinted platform allowed for integrating patient-derived cells and incorporating genetic variability early in drug development. The reduced potency in keratinocytes helps explain the limited benefit acyclovir and its congeners play in reducing sexual transmission. These data indicate that the 3D bioprinted HSE assay platform provides a more physiologically relevant approach to identifying potential antivirals for HSV.

**One Sentence Summary:** High-throughput screen using 3D bioprinted human skin equivalents to identify antivirals against HSV and evaluate cell-type specific effects.

## Introduction

Herpes simplex virus type 1 (HSV-1) and type 2 (HSV-2) cause recurrent oral and genital ulcer diseases and infect two-thirds of the global population (*1, 2*). Both HSV-1 and HSV-2 can cause neonatal herpes with high mortality and devastating neurological impairment (*3, 4*). HSV-2 infection increases the risk of human immunodeficiency virus (HIV) acquisition (5–7), with a recent study estimating 420,000 of 1.4 million newly acquired HIV infections annually attributable to HSV-2 infection (8). Despite successes in animal studies, candidate HSV-2 vaccines faltered in human trials (9–11). Acyclovir, the primary treatment for HSV infection, alleviates disease severity and shortens outbreak duration (12) but faces drug resistance in immunocompromised patients (13, 14), has limited efficacy in preventing subclinical reactivation, (15) and fails to address the increased risk of HIV transmission (16, 17). Thus, novel therapeutic strategies and antiviral agents are needed to address the public health burden of HSV infection.

Drug development relies heavily on 2D monolayers and animal models to assess drug safety and efficacy. Conventional *in vitro* cultured monolayers facilitate rapid and robust infectivity assays but don’t recapitulate complex cell-cell and cell-matrix interactions reflective of native tissues (18, 19). Consequently, these cellular assays often fail to predict drug responses in humans accurately (20). Animal models lack the predictive value and biological relevance to humans for effective drug discovery and development (21). This dilemma is evident in drug candidates failing to progress from Phase I to Phase III clinical trials, resulting in continued high costs for developing new pharmaceuticals (22, 23). 3D *in vitro* models closely mimicking human tissues and organs offer opportunities to circumvent the limitations in clinical predictability of current drug R&D platforms (24, 25).

Bioengineered tissues bridge the gap between existing cellular and animal models and the complexity of human hosts and offer a powerful platform for expediting drug discovery, toxicity screening, and preclinical testing in a high-throughput, cost-effective manner. These *in vitro* human tissue models are designed to replicate key features of *in vivo* organs, such as cell type composition, 3D microarchitecture, functional tissue interfaces, and organ-specific microenvironments (26). They offer unique opportunities for real-time visualization and high-resolution analysis of biological processes often unattainable in simpler cellular and animal models.

Skin and mucosal barriers are the primary tissue of concern for HSV initial infection and recurrent viral replication following reactivation. Skin tissue comprises the epidermis, dermis, and subcutaneous layer, forming a physical barrier protecting against pathogens and harmful environmental substances (27). The epidermis primarily consists of keratinocytes, which can differentiate into stratified layers, from the innermost stratum basale to the outmost stratum corneum. Human biopsy studies of recurrent HSV infection indicate basal keratinocytes at the dermal-epidermal junction (DEJ) serve as the replication targets for HSV reactivation through sensory nerve innervation and neurite extension into DEJ (28). Our recently developed 3D skin-on-chip also demonstrated that basal keratinocytes were the most susceptible targets for HSV infection during keratinocyte differentiation (29). Thus, both *in vivo* and *in vitro* evidence highlight the importance of using multicellular tissue models that recapitulate skin architecture when developing HSV therapeutics.

We developed bioprinted human skin equivalents (HSE) that differentiate into a stratified epidermis at the air-liquid interface (ALI) (30, 31). Here, we implemented a high-throughput screen (HTS) using 3D bioprinted HSE in a 96-well format and leveraging HSV-GFP reporter virus with high-content imaging. Our 3D bioprinted assay platform identified potent antiviral compounds, including 23 known anti-HSV candidates. We selected 11 compounds to evaluate in adult human skin-derived keratinocytes and fibroblasts, uncovering pharmacological properties distinct among cellular types in 2D and 3D, thus highlighting the importance of applying physiologically relevant assays in drug discovery and development.

## RESULTS

### Assessment of HSV Susceptibility and Acyclovir Antiviral Potency Across Different Cell Types

Herpes simplex virus enters the human body through the skin and mucosa surface and establishes latent infection for the lifetime of the host (32). The virus reactivates periodically and causes recurrent disease back to the periphery. Basal keratinocytes are the primary cells encountered by HSV reactivation in the skin (28, 29). However, current drug discovery practice utilizes Vero cell and fibroblast cultures for antiviral drug development (33). To investigate if acyclovir displayed similar effectiveness in keratinocytes, fibroblasts, and Vero cells, we isolated keratinocytes and fibroblasts from 6 donors (table S1) using 3mm punch skin biopsies (Fig. 1A). Real-time monitoring of recombinant strain HSV-1 K26 infection via GFP fluorescence (fig. S1) revealed GFP expression peaked at 20, 48, and 36 hour post-infection (HPI) in keratinocytes, fibroblasts, and Vero cells, respectively (Fig. 1B&C). Time to initial GFP detection was significantly earlier in keratinocytes (6.0 HPI) than in fibroblasts (7.9 HPI) (P < 0.001) (fig. S2A). GFP expression increased much faster in keratinocytes compared to both fibroblasts and Vero cells (P < 0.01), the maximum GFP doubling rate (mean ± sd) at 2.23 ± 0.41, 1.37 ± 0.39, and 1.00 ± 0.15 per hour in keratinocytes, fibroblasts, and Vero cells respectively (fig. S2B). These results indicated that HSV infection and viral gene expression occurred more rapidly in keratinocytes than in donor-matched fibroblasts or Vero cells.

**Fig. 1.**
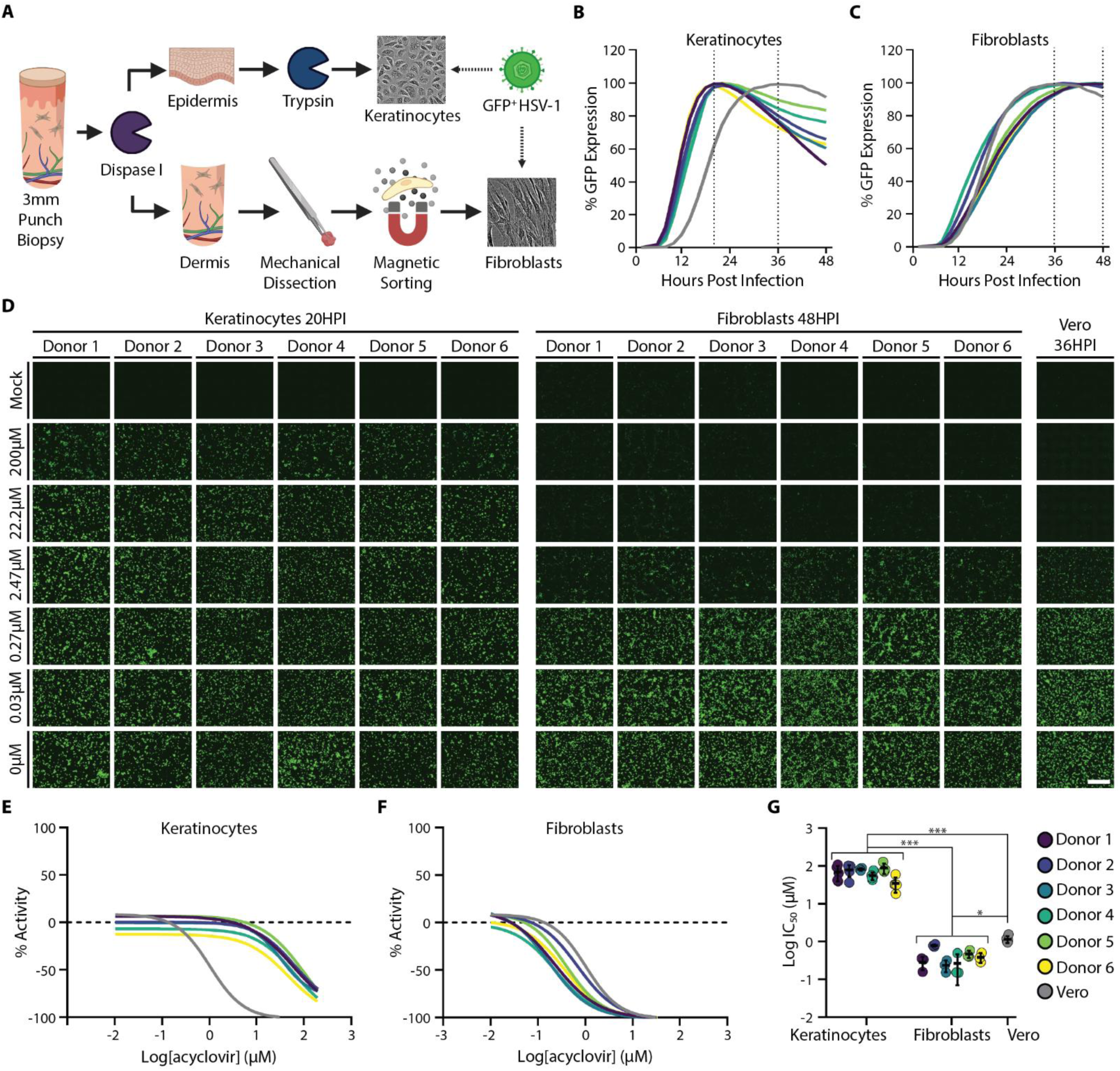
Acyclovir potency in donor-derived primary keratinocytes and fibroblasts and in Vero cells. **(A)** Punch biopsies from six donors were collected and dissociated by enzymatic and mechanical processes. Vero cells, keratinocytes **(B),** and fibroblasts **(C)** were infected with GFP-expressing HSV-1, and live cell images were taken every two hours. **(D)** Keratinocytes, fibroblasts, and Vero cells were infected with GFP-expressing HSV-1 and then treated with acyclovir at the specified doses. Representative live cell images were taken at the peak of GFP expression. (Scale bar 500µm). **(E)** Dose-response curve of acyclovir in keratinocyte cultures compared to Vero cells (grey line). **(F)** Dose-response curve of acyclovir in fibroblast cultures compared to Vero cells (grey line). **(G)** IC_50_ values for each donor in each cell type (*** *P* < 0.001, * *P* < 0.05, linear mixed model). Data represents averages of six donors; keratinocytes (N = 5 per donor), fibroblasts (N = 3 per donor), and Vero (N = 5) respectively. Error bars represent standard deviation.

We then evaluated the potency and efficacy of acyclovir in dose response using donor-matched keratinocytes and fibroblasts (Fig. 1D). Dose-response curves were established using the peak infection time previously determined in each cell type (Fig. 1E and F, and fig. S3). Acyclovir exhibited significantly reduced potency in keratinocytes compared to fibroblasts across all six donors (Fig. 1D-G, *P* < 0.001). The concentration of acyclovir required to inhibit 50% of virus-encoded GFP expression (IC_50_) for each donor was, on average (mean ± sd), 196.7-fold higher in keratinocytes (67.7± 18.2μM) than in fibroblasts (0.40 ± 0.2μM) and 60.2-fold higher than in Vero cells (1.14 ± 0.2μM). The IC_50_ of acyclovir in keratinocytes was also over two-fold higher than published peak serum levels in patients following treatment with three times daily 1000mg valacyclovir (34). This suggests that current acyclovir treatment might not be optimal for inhibiting HSV gene expression in the skin epidermis. These results confirmed that antiviral drug potency can vary substantially among cell types tested, emphasizing the importance of employing physiologically relevant cells in antiviral drug testing.

### Development of HSV Infection Assays on 3D Bioprinted Human Skin Equivalents

The significant differences in acyclovir responses between keratinocytes and fibroblasts led us to develop multicellular tissues for early drug discovery that better recapitulate skin *in vivo* to investigate antiviral potency and efficacy. To create physiologically relevant *in vitro* human skin models of HSV-1 infection for antiviral screening, we 3D bioprinted human skin equivalents (HSE) in a 96-transwell plate format. Using a hydrogel containing gelatin, collagen, fibrin, and neonatal human fibroblasts, the dermal tissue was bioprinted onto the apical side of a 96 transwell insert using a plunger-based 3D bioprinter (Fig. 2A). The dermal tissues underwent 7 days of maturation before neonatal human keratinocytes were seeded on top and submerged in epidermalization media for 7 more days. To induce keratinocyte differentiation and epidermis stratification, the tissues were lifted to air-liquid interface (ALI) using a custom 3D printed adaptor (fig. S4) and cultured in cornification media at the basal surface of the dermis for 7 days (Fig. 2A) (30, 31, 35). Hematoxylin and eosin (H&E) staining confirmed that ALI cultures produced HSE with a dermis and differentiated epidermis, including a stratum basale and stratum corneum (Fig. 2B column 1). Immunohistochemistry (IHC) staining demonstrated cytokeratin 10 (K10) and cytokeratin 14 (K14) expression in the suprabasal and basal layer, respectively, indicating properly differentiated epidermis resembling *in vivo* human skin architecture (Fig. 2B column 2) (36).

**Fig. 2.**
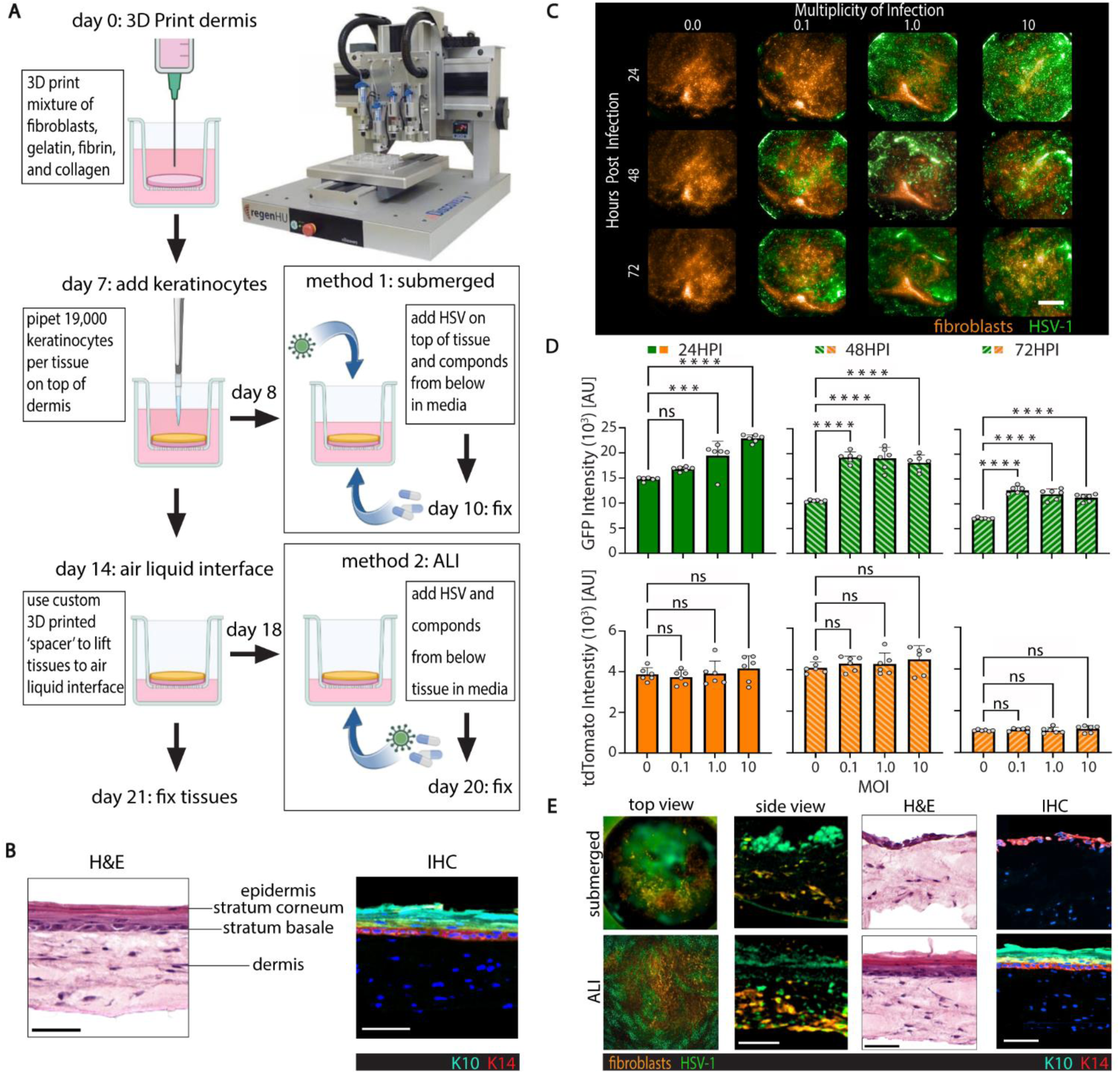
3D bioprinted HSE assay development and validation. **(A)** Dermis equivalents were 3D printed onto the apical side of transwell inserts using the RegenHU 3D Discovery bioprinter (image courtesy of RegenHU). Keratinocytes were pipetted onto the apical surface of the dermis. In the submerged model, the tissues were infected at the apical surface. In the ALI model, tissues were brought to ALI and then infected at the basolateral surface (created with BioRender.com). **(B)** H&E and IHC images of differentiated ALI tissues. K10 (cyan) and K14 (red) identify keratinocytes in the suprabasal and basal layer of the epidermis respectively (scale bar 50µm) **(C)** Submerged tissues were infected at various MOI and then imaged at specified times. Fibroblasts express tdTomato (orange) while infected cells express GFP (green) (scale bar 1mm). **(D)** GFP and tdTomato signal at each MOI and timepoint (N = 6, *** *P* < 0.001, **** *P* < 0.0001 by ordinary one-way ANOVA). **(E)** Maximum projection of infected tissues from the top (column 1) or side (column 2) view. H&E (column 3) and IHC (column 4) staining of infected submerged or ALI models (scale bar 1mm (column 1) or 50µm (column 2, 3, 4).

The antiviral screen was completed using two fluorescent markers; GFP-fused HSV-1 strain K26 (37) was employed to measure virus infection, while fibroblasts were transduced to constitutively express tdTomato to monitor compound cytotoxicity. The infection conditions were optimized by imaging the tissues using a high-content microscope at 24, 48, and 72 hours post-infection (HPI). The total fluorescence signal of GFP and tdTomato was quantitated using a maximum projection at a multiplicity of infection (MOI) of 0.1, 1.0, and 10 PFU/cell (Fig. 2C). GFP expression had the most significant difference between mock and infected using an MOI of 0.1 at 48HPI (difference in relative fluorescence units (*ΔRFU*) = 8753, *P* < 0.001), while the tdTomato signal was not affected (*ΔRFU* = 201.9, *P* = 0.8828) by viral infection (Fig. 2D). We applied these optimal infection conditions to subsequent experiments.

We used two different infection methods, submerged and ALI cultures, to emulate infection routes for primary and recurrent viral encounters. The submerged culture maintained an undifferentiated single epithelium layer and was infected apically, resembling HSV primary infection at initial viral exposure. The ALI model included a stratified epidermis infected basolaterally, mimicking viral reactivation from the dermis. In the submerged model, apical infection of HSV-1 resulted in GFP expression mainly detected in keratinocytes above the tdTomato-positive dermal fibroblasts (Fig 2E, top row, column 2). In ALI cultures infected from the basolateral route of infection, there was colocalization of GFP and tdTomato signals, suggesting infection mainly occurred in the fibroblasts (Fig. 2E, bottom row, column 2). H&E and IHC staining demonstrated that infected submerged cultures had a disrupted epithelial monolayer, while infected ALI tissues maintained stratified epidermal morphology with proper K10 and K14 expression (Fig. 2E, columns 3 and 4). Since the submerged and ALI infection models preferentially targeted keratinocytes and fibroblasts, respectively, we used both models for anti-HSV drug screening.

To validate our screening assay pharmacologically, we tested acyclovir in dose-response in submerged and ALI models (fig. S5). The IC_50_ of acyclovir was > 9-fold higher in the submerged tissues (IC_50_ = 0.36μM), which infect mainly keratinocytes, compared to the ALI skin tissues (IC_50_ < 0.04μM), which infect predominantly fibroblasts. As expected, acyclovir inhibited HSV-1 gene expression with minimal cytotoxic effects on the fibroblasts. These data confirmed that our 3D HSE model could capture cell-type specific antiviral properties of acyclovir.

### Implementation of a Primary Drug Screen using 3D Bioprinted Human Skin Tissue Equivalents

We implemented a screen of 738 compounds with broad targets and a wide range of mechanisms of action (MOA) (spreadsheet S1) utilizing our 3D bioprinted HSE assay platform. A primary screen tested the compound library in both the submerged and ALI infection models at 10μM compound concentration in duplicate, using 3,840 bioprinted HSE tissues. Each plate contained control wells: negative control (HSV-1 + DMSO), inhibitor control (media -HSV-1), and a known compound (HSV-1 + ACV (0.2μM)) (Fig. 3A). We used the maximum projection of each well to determine the total fluorescence of GFP (HSV-1 activity) and tdTomato (fibroblast viability) and normalized the data to the control wells in the corresponding test plate as described in Materials and Methods (Fig. 3B, fig. S6, spreadsheet S2).

**Fig. 3.**
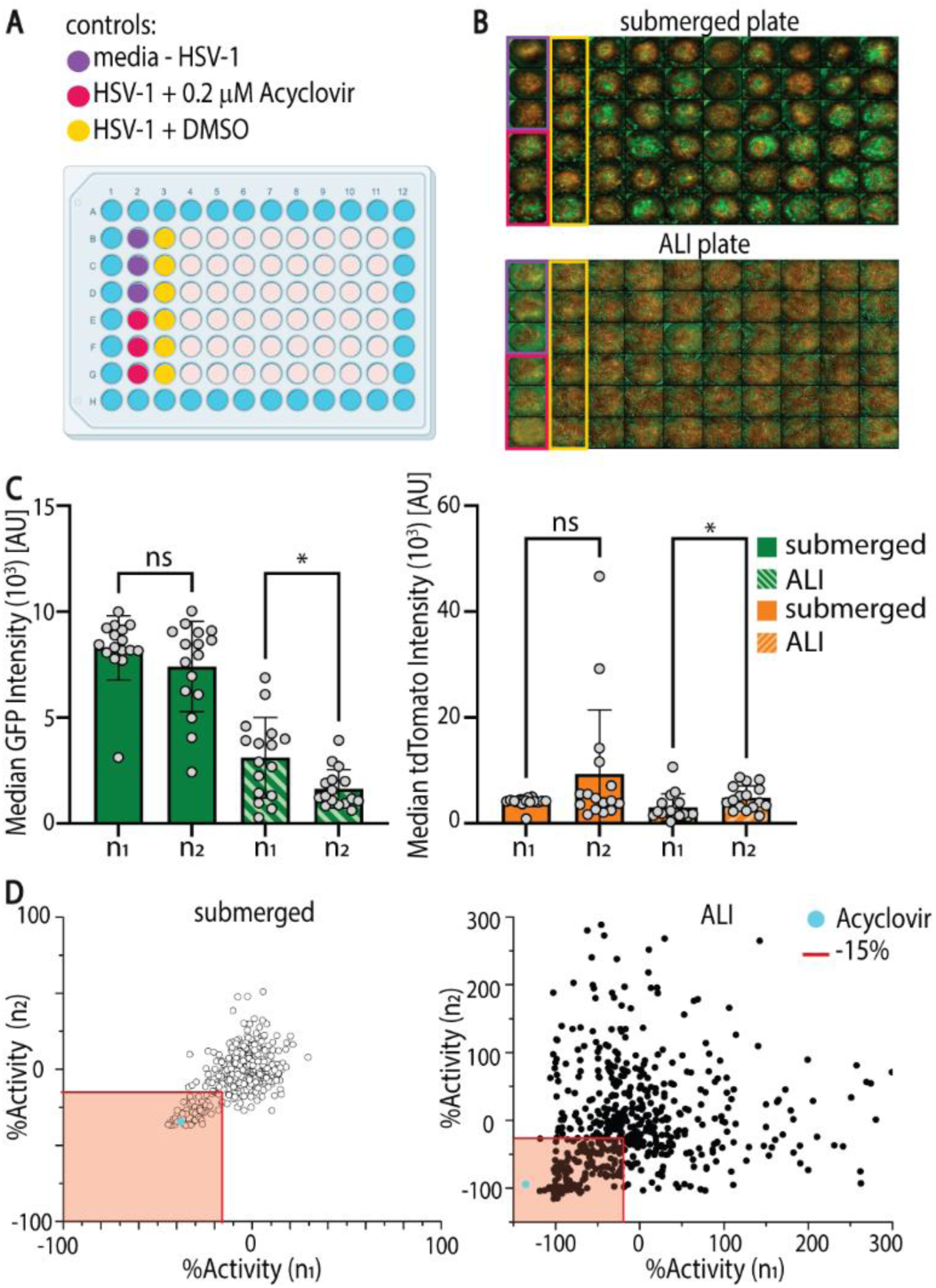
Primary screen of compound library in 3D bioprinted assay platform. **(A)** Plate layout for compound screen. **(B)** Representative images from primary screen of submerged and ALI models. Controls are highlighted in boxes with colors corresponding to part (A). **(C)** Wilcoxon two-sample analysis of GFP intensity (left) and tdTomato intensity (right) of the median of the six negative control wells (HSV-1 + DMSO) across two replicate plates (N =16, * *P* < 0.05). Bars represent average with error bars as standard deviation. **(D)** Correlation plots of the two replicates for the primary screen of compounds. The red shaded box represents compounds that were selected as hits for re-testing if they reduced GFP expression (%Activity) by at least 15% in either replicate or model (left submerged, right ALI). Acyclovir was included in the collection and denoted by a cyan circle.

To confirm the reproducibility of our assay, we compared the median GFP and tdTomato fluorescence signal from the negative control wells between the two assay repeats (*n_1_* vs *n_2_*) in both the submerged and ALI models. The difference in the GFP or tdTomato intensity of the control wells was not significant in submerged models, indicating that our HSV-1 gene expression and fibroblast viability were consistent among all 32 plates (Fig. 3C, *P_GFP_* = 0.3045, *P_tdTomato_* = 0.5641). The difference in the GFP and tdTomato intensity of the control wells in the ALI plates was significant (Fig. 3C, *P_GFP_* = 0.0234, *P_tdTomato_* = 0.0121), emphasizing the need to include and normalize to controls on each plate to correct for any variability.

To determine if our assay is amenable to high-throughput screening (HTS), we calculated the median *Z’*-factor, a measure of the robustness of the assay. A score that is Z’>0.5 denotes a robust assay window for screening, 0.5>Z’>0 indicates a marginal assay window and that the screen needs replicates, and a Z’<0 means that the assay window is not robust enough for screening. In the submerged model, the Z*’* was 0.57, and for the ALI model, the *Z’* was 0.13, indicating that the submerged assay has excellent robustness, while the ALI assay has moderate robustness, demonstrating that our 3D bioprinted assay platform was amenable to HTS (fig. S7) (38). We identified ‘hits’ as compounds that inhibited GFP expression (HSV-1 activity) by at least 15% without causing more than 50% reduction in tdTomato fluorescence (fibroblast viability) in either screen mode (Fig. 3D), selecting 106 compounds from the primary screen.

### Secondary Drug Testing with 3D Bioprinted Human Skin Equivalents

Next, we performed a secondary screen of the 106 selected ‘hits’ in dose-response from 10µM to 0.04µM in the submerged and ALI models (spreadsheet S3). We found that 46% of compounds in the submerged and 70% compounds in the ALI model reduced GFP expression (%Activity) by at least 50% at the maximum concentration tested without reducing tdTomato signal by 50% (%Viability) (Fig. 4A). Tested compounds were categorized by concentration-response curve (CRC) class (Fig. 4B) (38), according to their potency (IC_50_) and efficacy (maximum percent activity). Compounds were considered candidate antivirals if they had CRC classes of -1.n or -2.n for HSV-1 inhibition and CRC class of 4 for fibroblast viability in at least one infection model. Forty-one candidate antivirals were identified and grouped by MOA (Fig. 4C). The MOA clustering indicated that HDAC inhibitors (8%) and protease inhibitors (8%) were more prevalent candidates in the submerged model while DNA polymerase inhibitors (13%) were more prevalent antivirals in the ALI model (fig. S8). Inhibitors of proteasome, DNA polymerase, Casein Kinase 2, Exportin-1, GBF-1, and Ribonucleotide-Reductase were effective in both models. Of the 41 candidate antivirals, 23 are known or experimental HSV treatments (Fig. 4D), including five ‘ciclovir’ compound, the most common class of HSV antivirals (Fig. 4E, table S2) (39–59), demonstrating our assay’s ability to identify HSV antivirals in an unbiased screen.

**Fig. 4.**
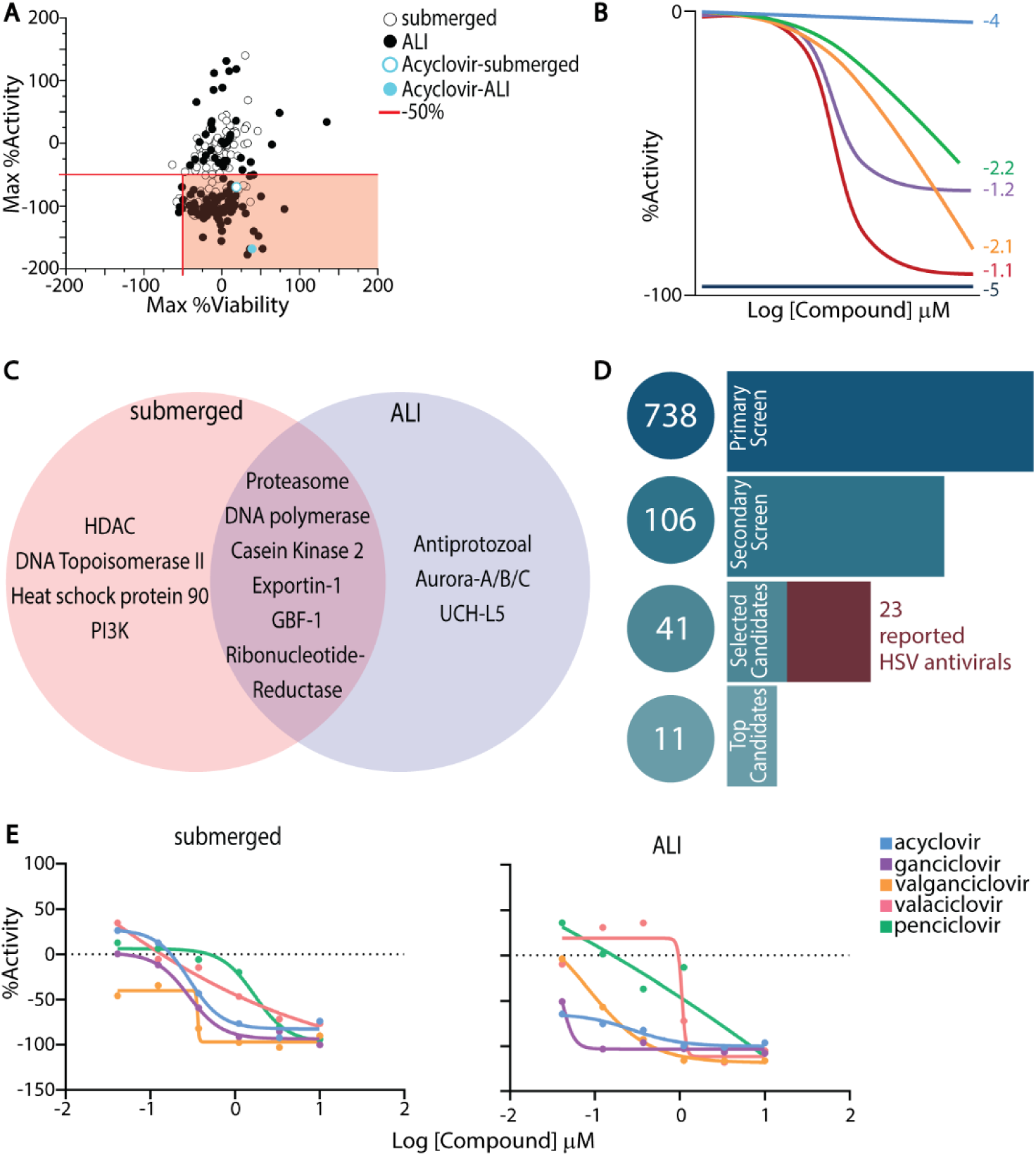
Dose response of candidate antivirals in 3D bioprinted assay platform. **(A)** Correlation plot of Max %Activity (maximum reduction in GFP) vs. Max %Viability (maximum reduction in tdTomato) of 106 ‘hits’ tested in dose response. Top candidate antivirals (50% or greater reduction in GFP) that did not kill over 50% of tdTomato transduced fibroblasts are identified by the red shaded box. **(B)** Schematic illustrating %Activity dose response profiles of different Concentration-Response Curve classes (CRC). **(C)** Venn diagram showing divergent and coinciding targets for 41 top candidate antivirals in both submerged and ALI models. **(D)** Schematic of compounds selection from 738 compounds in the primary screen to 106 ‘hits’ tested in dose-response to 41 selected candidates and 11 top candidates selected to move forward. Of the 41 selected candidates, 23 are current or experimental HSV treatments. **(E)** Dose response curves of candidate antivirals in “ciclovir” family, known to treat HSV-1, in submerged and ALI models (N = 1).

Based on their CRC classification, antiviral potency, and low fibroblast toxicity, we selected 11 of the 41 compounds as top-candidate antivirals for further testing. These included two known anti-herpes drugs, amenamevir and pritelivir (41, 45, 56), and four that have been reported to have anti-herpes activity: gemcitabine (51), lanatoside-C (53), niclosamide (47), and SNX-2112 (57). We re-tested these compounds in triplicate to assess potency and efficacy (Table 1, Fig. 5A). The IC_50_ and maximum inhibition (Fig. 5B and C) indicate that compounds were generally more potent in the ALI than the submerged model in 10 of the 11 compounds tested (Table 1). Three compounds, fimepinostat, LDC4297, and VLX1570, showed significant differences in potency between the two models (Fig. 5B). We next plotted the viability of the tdTomato fibroblasts (Fig. 5D) and calculated the cytotoxic concentration required to kill 50% of target cells (CC_50_) (Table 1). While none of the compounds were cytotoxic in the ALI model, lanatoside-c, SNX-2112, and VLX1570 exhibited some cytotoxicity in submerged models at higher concentrations.

**Fig. 5.**
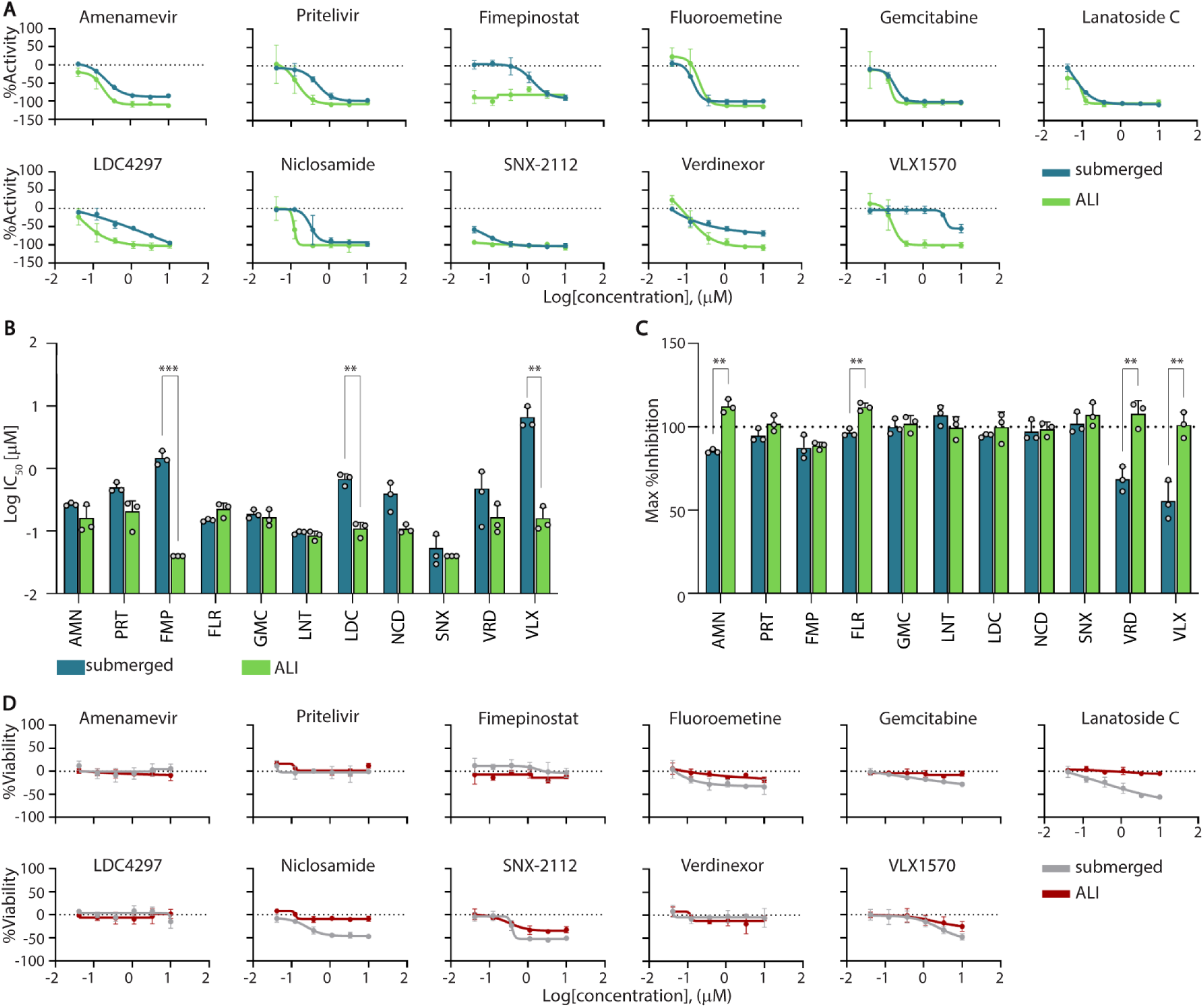
Top candidate antiviral potency, efficacy, and cytotoxicity in 3D bioprinted HSE. **(A)** Average dose-response curves for the 11 top candidate antivirals for submerged (blue) and ALI (green) models. **(B)** IC_50_ values for each top candidate antiviral compared between submerged (blue) and ALI (green) models. **(C)** Maximum inhibition for each top candidate antiviral compared between submerged (blue) and ALI (green) models. Statistical significance was determined by linear mixed model (*** *P* < 0.001, ** *P* < 0.01, * *P* < 0.05) for (B) and (C). **(D)** CC_50_ dose response curves for 11 top candidate antivirals. All data is plotted as the average of three replicates (N = 3), error bars represent standard deviation.

**Table 1:** Pharmacologic profile of top candidate antivirals. Top candidate antivirals were selected from 41 compounds identified as potent and effective at doses below their respective CC_50_ in the primary and secondary screens. We re-tested these top candidate antivirals in triplicate and reported the Curve Class, Max Inhibition, IC_50_, and CC_50_. Known anti-HSV compounds, amenamevir and pritelivir, highlighted in grey.

| Candidate antiviral | Primary Mechanism of Action | Target | submerged |  |  |  | ALI |  |  |  |
| --- | --- | --- | --- | --- | --- | --- | --- | --- | --- | --- |
|  |  |  | Curve Class | Max Inhibition (%) | IC<sub>50</sub> (μM) | CC<sub>50</sub> (μM) | Curve Class | Max Inhibition (%) | IC<sub>50</sub> (μM) | CC<sub>50</sub> (μM) |
| Amenamevir (AMN) | Helicase primase inhibitor | Viral | -1.1 | 85.31 | 0.27 | > 10 | -1.1 | 112.36 | 0.16 | > 10 |
| Pritelivir (PRT) | Helicase primase inhibitor | Viral | -1.1 | 94.64 | 0.50 | > 10 | -1.2 | 101.90 | 0.21 | > 10 |
| Fimepinostat (FMP) | PI3K/HDAC inhibitor | Host | -1.1 | 87.43 | 1.48 | > 10 | -5 | 88.60 | < 0.04 | > 10 |
| Fluoroemetine (FLR) | Unknown | Host | -1.1 | 96.50 | 0.15 | > 10 | -1.1 | 111.83 | 0.22 | > 10 |
| Gemcitabine (GMC) | Ribonucleotide Reductase | Host | -1.1 | 99.84 | 0.19 | > 10 | -1.2 | 101.78 | 0.16 | > 10 |
| Lanatoside C (LNT) | Autophagy inducer | Host | -1.1 | 106.86 | 0.09 | 2.49 | -1.2 | 99.49 | 0.08 | > 10 |
| LDC4297 (LDC) | CDK inhibitor | Host | -2.1 | 94.82 | 0.68 | > 10 | -1.1 | 99.87 | 0.11 | > 10 |
| Niclosamide (NCD) | Multi-functional | Host | -1.1 | 97.13 | 0.39 | > 10 | -1.1 | 98.67 | 0.11 | > 10 |
| SNX-2112 (SNX) | HSP90 inhibitor | Host | -1.2 | 101.78 | 0.05 | 1.14 | -5 | 107.14 | < 0.04 | > 10 |
| Verdinexor (VRD) | Exportin Antagonist | Host | -1.3 | 68.57 | 0.48 | > 10 | -1.1 | 107.81 | 0.17 | > 10 |
| VLX1570 (VLX) | Protease deubiquitinase inhibitor | Host | -2.5 | 55.49 | 6.67 | 8.41 | -1.1 | 101.22 | 0.16 | > 10 |

### Assessment of the Candidate Antiviral Activities in Donor-Derived Keratinocytes and Fibroblasts

Differences in the IC_50_ values between the submerged and ALI models suggested that, like acyclovir, some drugs may exhibit cell-type specific antiviral properties. We next investigated the 11 top candidate antivirals in our 2D monocultures of donor-derived primary keratinocytes and fibroblasts in dose-response to further investigate cell-type specific effects, in comparison to acyclovir (Fig. 6A). According to the IC_50_ and maximum inhibition, all 11 candidate antivirals were more potent than acyclovir in keratinocytes (7 to >884-fold), with IC_50_ values ranging from 0.08 to 9.9μM (Fig. 6, A, B, and C, and fig. S9). Five candidate antivirals, amenamevir, pritelivir, gemcitabine, lanatoside-C, and SNX-2112, surpassed acyclovir antiviral potency in fibroblasts as well, with IC_50_ values ranging from 0.05 to 0.15μM (Fig. 6A and B, fig. S10).

**Fig. 6.**
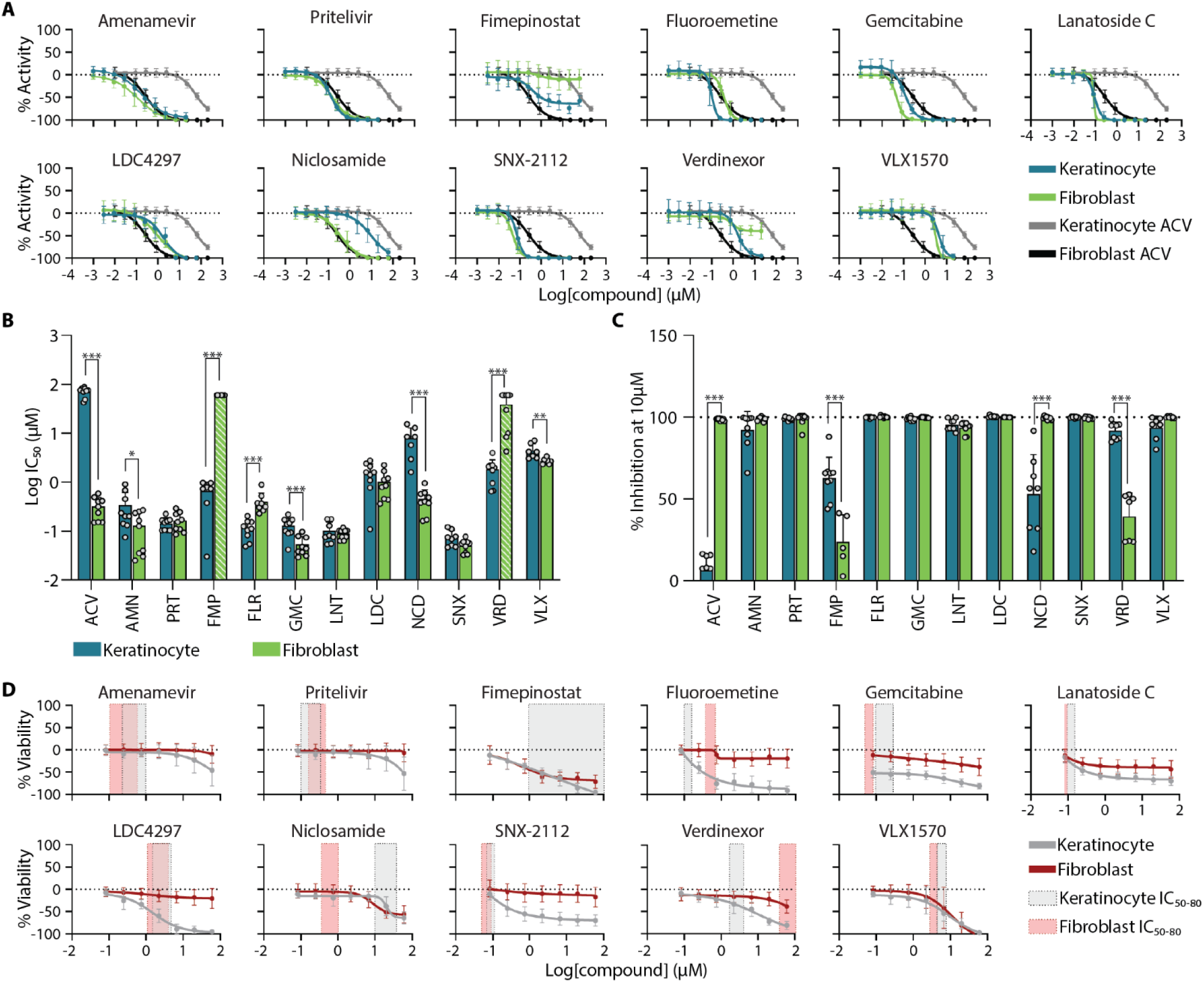
Candidate antiviral potency, efficacy, and cytotoxicity in donor-derived primary keratinocyte and fibroblast monocultures. **(A)** Dose-response curves for the 11 top candidate antivirals compared to acyclovir (ACV) (keratinocytes blue and grey, respectively; fibroblasts green and black, respectively). **(B)** IC_50_ values for each top candidate antiviral compared between keratinocytes (blue) and fibroblasts (green). Striped bars (FMP, VRD) indicate candidate antivirals that failed to reduce GFP expression by at least 50% consistently. **(C)** Maximum inhibition for each top candidate antiviral is compared between keratinocytes (blue) and fibroblasts (green). Statistical significance was determined by linear mixed model (*** *P* < 0.001, ** *P* < 0.01, * *P* < 0.05) for (B) and (C). **(D)** CC_50_ dose-response curves for all twelve candidate antivirals compared to their respective IC_50_ to IC_80_ dose ranges. Keratinocyte data is from 20HPI (grey), while fibroblast data is from 48HPI (red). All data is plotted as the average of three replicates in three distinct donors (N = 9), error bars represent standard deviation.

Antiviral potency differed significantly between keratinocytes and fibroblasts for 8 of the 12 compounds in donor-matched skin cells (Fig. 6B and C, fig. S11). However, the differences in candidate antivirals were less than in acyclovir (<27 fold versus >220 fold, respectively). Two notable exceptions were verdinexor and fimepinostat, which inhibited viral GFP expression by at least 50% in keratinocytes but failed to inhibit viral GFP expression in fibroblasts reliably. We also detected significant differences in antiviral responses between donors 3 and 4 (P < 0.001) and donors 5 and 4 (P = 0.02) when considering potencies for all 11 compounds in keratinocytes, but not fibroblasts (fig. S12-14, table S3).

We next examined compound toxicity on keratinocytes and fibroblasts in the absence of viral infection (Fig. 6D, fig. S9&10). Keratinocytes were generally more sensitive to compound cytotoxicity than fibroblasts. Amenamevir and pritelivir, which target viral helicase/primase, showed a highly effective antiviral dosage (IC_50_ through IC_80_), which was at >100-fold range lower than the CC_50,_ yielding a selectivity index of >250 in keratinocytes and >400 in fibroblasts (Fig. 6D, table S4). On the other hand, gemcitabine, SNX-2112, and lanatoside-C, which inhibit host pathways involving DNA replication, have high selectivity indexes (>750-1200) only in the fibroblasts and showed toxicity in keratinocytes.

To confirm candidate antivirals were effective against HSV-2 as well, we conducted a plaque reduction assay. Primary keratinocytes in 6-well plates were infected with either HSV-1 K26 or HSV-2 186 and treated with multiple doses of each candidate antiviral. All 11 candidate antivirals inhibited HSV-1 and HSV-2 plaque formation at similar doses (fig. S15).

### Comparison of Candidate Antivirals in 2D Monoculture versus 3D Bioprinted Human Skin Equivalents

We compared the potency, toxicity, and selectivity index of the top 11 candidate compounds in our four models: 2D keratinocytes, 2D fibroblasts, 3D submerged, and 3D ALI (Fig. 7, A and B, table S5). The impact of cell type and 2D:3D model type on the antiviral potency was compound-specific. Acyclovir was most significantly affected by cell type and the 2D:3D environment, whereas lanatoside C and SNX-2112 were unaffected (table S6). Submerged tissues and keratinocytes displayed more similar potencies than ALI tissues and fibroblasts, most likely due to the undifferentiated monolayer of keratinocytes infected in the submerged model (Fig. 7, C and D). Combining cell type differences between keratinocytes and fibroblasts and the 2D:3D growth environment, we showed that amenamevir, pritelivir, fluoroemetine, gemcitabine, lanatoside C, and SNX-2112 exhibited less than a 3-fold difference in average potency between models (Fig. 7C-E); all remaining candidate antivirals were >3-fold more potent in the 3D model. Keratinocyte 2D monocultures were more susceptible to cytotoxic effects of candidate antivirals than 2D monocultures of fibroblasts or the 3D models, which identify toxicity through a loss of fluorescently labeled fibroblasts (Fig. 7B). All candidate compounds were selective (selectivity index >10) in the ALI model, and 9 of the 11 were selective in submerged models. While 8 of the 11 candidate antivirals were selective in 2D fibroblasts, only the two helicase-primase inhibitors were selective in 2D keratinocytes. Antiviral potency and toxicity can significantly differ between cell type and 2D:3D model tested, emphasizing the importance of applying physiologically relevant models to assess drug responses early in development.

**Fig 7:**
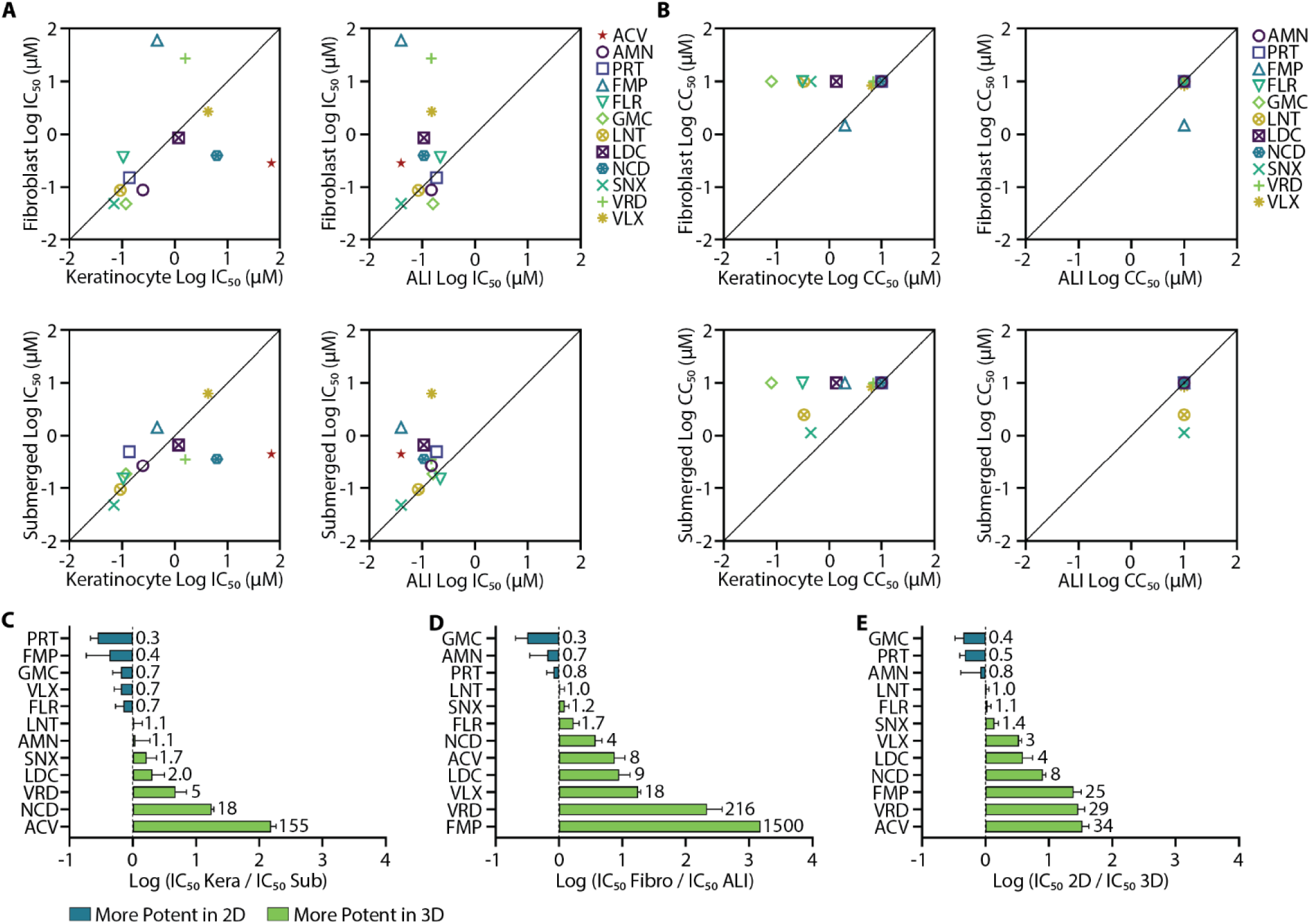
Comparison of 3D and 2D models in testing novel antiviral candidates. **(A)** Pairwise comparisons of IC_50_ values for candidate antivirals in the four models tested. **(B)** Pairwise comparisons of CC_50_ values for candidate antivirals in the four models tested. **(C)** Fold change was determined by dividing the IC_50_ value of each candidate antiviral in keratinocytes by the IC_50_ of the same candidate antiviral in submerged models. **(D)** Fold change was determined by dividing the IC_50_ value of each candidate antiviral in fibroblasts by the IC_50_ of the same candidate antiviral in ALI models. **(E)** IC_50_ values for each candidate antiviral were pooled (keratinocytes and fibroblasts, submerged and ALI), then IC_50_ values for candidate antivirals in 2D were divided by IC_50_ values in 3D. (C) (D) (E) Green bars indicate candidate antivirals that were more potent in 3D, while blue bars indicate candidate antivirals that are potent in 2D. Data for 2D (N = 9) and 3D (N = 3) is reported as averages, error bars are standard deviation.

### Generation of 3D Bioprinted Human Skin Equivalents using Adult Donor-Derived Keratinocytes

Human neonatal skin cells are readily available commercially, but the ability to use adult donor-derived cells would increase the translational impact of our drug discovery assay platform. We bioprinted the dermis with commercial neonatal human dermal fibroblasts, then seeded the dermis with Donor 3 keratinocytes as a proof of concept to generate donor-specific HSE. H&E staining of submerged and ALI HSE showed donor-derived keratinocytes successfully developed into a stratified epidermis in the adult-derived ALI model with proper expression of K10 and K14 determined by IHC (Fig 8, A and B). Most of the 11 candidate antivirals displayed similar IC_50_ values in neonatal-derived and adult donor-derived HSE (Fig. 8, C, and D, table S7). Thus, our assay platform could use easily accessible neonatal cells to initially screen large numbers of compounds inexpensively and then incorporate adult human skin cells for further preclinical evaluation to enhance drug testing in diverse genetic backgrounds.

**Fig 8:**
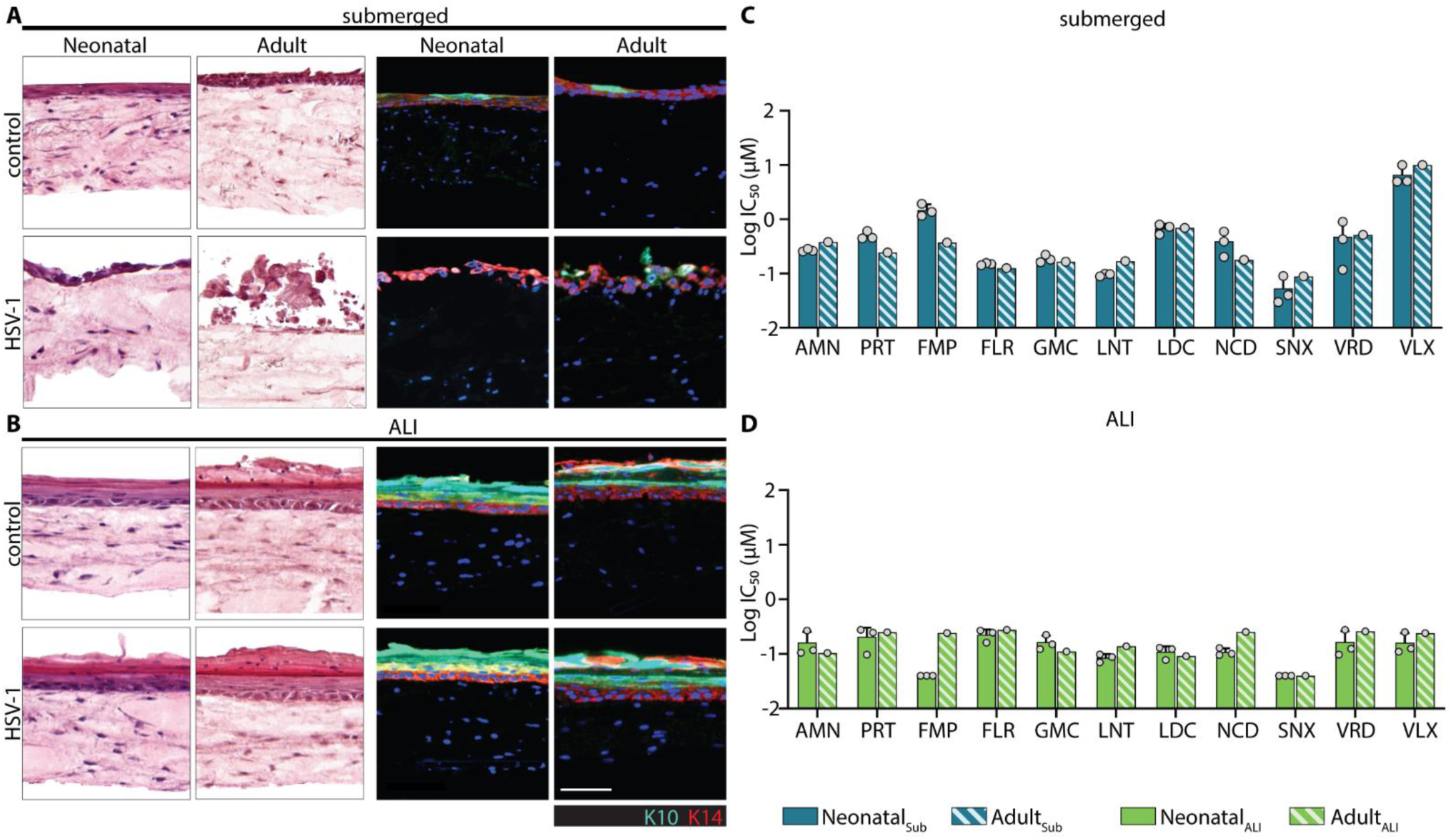
Generation of 3D bioprinted HSE using adult donor-derived keratinocytes. H&E and IHC staining of commercial neonatal or adult-derived bioprinted HSE in submerged **(A)** and ALI **(B)** models. Comparison of calculated IC_50_ in submerged **(C)** and ALI **(D)** models for top candidate antivirals. Neonatal data is plotted as an average of three replicates (N = 3), error bars are standard deviation. Adult data has an N = 1.

## DISCUSSION

Our study highlights critical variations in the susceptibility of different cell types to HSV manifestation and the effectiveness of acyclovir, a widely used antiviral drug for treating herpes diseases. We have demonstrated that acyclovir exhibits substantially lower potency in suppressing HSV infection in keratinocytes than in fibroblasts or Vero cells. Keratinocytes are the primary cell type encountered during HSV reactivation and peripheral infection (28, 32). The faster viral replication observed in these cells suggests keratinocytes are the more physiologically relevant targets for antiviral drug screening and preclinical testing. This underscores an important limitation of current antiviral testing approaches, which utilize Vero cells and fibroblasts monolayer cultures, and has significant implications for HSV drug development strategies. We have established a novel high-throughput assay platform leveraging 3D bioprinting to rapidly generate over 6,500 human skin equivalents (HSE) consisting of keratinocytes and fibroblasts to identify potent HSV antiviral compounds. The 3D bioprinted HSE recapitulates the *in* vivo skin architecture, including the epidermis, dermis, and dermal-epidermal junction (DEJ), where reactivating HSV is released into the periphery (28). A GFP-expressing HSV-1 strain and tdTomato-expressing fibroblasts allowed our platform to rapidly screen for potency, efficacy, and cytotoxicity using 738 distinct compounds. We identified 41 potential antiviral candidates, 23 of which are known or experimental HSV antivirals, demonstrating the ability of our assay to identify drugs active against HSV in an unbiased screen. Finally, the successful integration of donor-derived adult skin cells into our platform to test antiviral responses further enhances the translational relevance of our preclinical drug screening system.

We demonstrated that the potency and efficacy of antiviral compounds could vary significantly between cell types and 2D versus 3D environments. The varied drug potency and efficacy between donor-matched keratinocytes and fibroblasts is best illustrated by acyclovir. This mainstay drug suppresses HSV lesion formation and reduces the duration of ulcer lesions but has limited efficacy in preventing subclinical shedding and transmission to sexual partners (12, 15). The IC_50_ of acyclovir in donor-derived keratinocytes was over twice the peak of clinical serum levels following three times daily treatment with 1000 mg valacyclovir (34), a dose substantially higher than 1000mg once per day used for standard suppressive therapy with valacyclovir (60). Furthermore, mathematical modeling predicts that current treatment regimens result in ineffective doses of acyclovir for over 40% of the day (61). The higher IC_50_ of acyclovir in keratinocytes we describe here and the relatively short half-life of acyclovir (62) likely contribute to the episodic cycle of asymptomatic shedding during suppressive therapy. Together, the increased HSV gene expression and reduced acyclovir potency in keratinocytes might explain why acyclovir therapy cannot eliminate subclinical shedding, the associated risk of HSV transmission, and the increased risk of HIV acquisition (15, 16, 63).

Our multi-model drug screening and evaluation strategy also yielded new insights into HSV antiviral drug development. Our data showed antivirals like amenamevir and pritelivir (41, 56), which inhibit HSV helicase-primase, were highly selective in tested 2D and 3D models and exhibited similar potencies against HSV-1 and HSV-2 strains, in agreement with published work (41, 56). Drugs that exhibited less than a 2-fold difference in potency for all models included pritelivir, amenamevir, gemcitabine, SNX-2112, and lanatoside C. Interestingly, the first four of these drugs directly target viral DNA synthesis, while lanatoside C functions by preventing viral genetic material from entering the nucleus (41, 53, 56, 57, 64). While our data suggested that targeting viral helicase-primase machinery was safe and effective, drugs targeting host cell replication machinery for DNA synthesis could be effective across multiple cellular environments when toxicity is not observed in complex models (65).

The reduced efficacy of acyclovir in keratinocytes could reflect inherent biological differences in drug metabolism and viral replication dynamics between cell types. Acyclovir is an acyclic purine nucleoside analog and functions as a prodrug. It is activated through phosphorylation by HSV thymidine kinase. Following this initial phosphorylation, two more phosphates are added to ACV-P through cellular kinase, resulting in ACV-triphosphate to terminate DNA chains and eliminate HSV (66). Thus, for acyclovir to inhibit viral DNA synthesis, it must be processed by cellular thymidine kinase in two of the three phosphorylation steps, which could introduce a source of cell-type-specific variability. In keratinocytes, HSV expression occurred more rapidly with a shorter lag time and faster kinetics than in Vero cells or donor-matched fibroblasts, suggesting their greater susceptibility to HSV infection and replication, which might be related to stem-like cellular nature of basal keratinocytes. While targeting viral DNA synthesis was generally effective in all models tested, acyclovir was substantially less potent in keratinocytes than fibroblasts, as demonstrated in 2D monolayer cultures and 3D bioprinted HSE. Future research is needed to uncover these mechanisms to improve antiviral strategies.

There are several aspects in which our engineered skin tissue models can be improved for higher physiological relevance to HSV infection. First, we could improve infection of stratified epithelium in the ALI culture through wounding at the apical surface to bypass the physical barrier of cornified epithelium. Second, our 3D models measure off-target drug toxicity using the tdTomato signal expressed by fibroblasts. Future iterations of this model could incorporate a third fluorophore into keratinocytes to measure toxicity. Third, constructing HSV-2 GFP expressing recombinant virus similar to HSV-1 K26 would benefit more direct and rapid testing against HSV-2 infection. Other strategies of antiviral drug development against wildtype and clinical isolates of HSV could include modifying target cells to express GFP under a viral promoter.

Our assay platform can also be expanded by incorporating more donor-derived cells into the skin tissues. Incorporating broad genetic diversity for drug screening could improve clinical trials and patient outcomes when variable cellular factors heavily impact candidate drug efficacy (67, 68). Our platform’s ability to rapidly screen many different compounds could also be leveraged to screen combination therapy between a select group of compounds. Combination therapy could be used to overcome cell-type specific pharmaceutical properties or prevent the emergence of viral resistance, like the approach employed by antiretroviral therapy in treating HIV infection.

In summary, we have established a 3D bioprinted HSE assay platform that can be used for drug screening to identify anti-viral compounds in a high-throughput format. We demonstrated that the potency and efficacy of candidate antivirals can vary significantly between cell types in both 2D cellular and 3D tissue cultures, with substantially lower potency observed for acyclovir in keratinocytes. Further, we identify that cell-type specific differences are largely absent in helicase-primase inhibitors, supporting the continued investigation of both this class of antivirals and novel antivirals targeting viral proteins. Finally, we establish that our assay platform can incorporate donor-derived cells, allowing early stages of drug discovery to include multiple donors with varied ages, sex, ancestry, and genetic backgrounds to improve success in downstream clinical trials.

## METHODS

### Study Design

3D human skin equivalents (HSE) were generated in controlled experiments. Commercial neonatal fibroblasts and keratinocytes were expanded and frozen so that every experiment was performed at the same passage and using the same lot numbers. HSV-1 stocks were stored at -150°C and only used once to avoid freeze-thaw cycles. Candidate antivirals were spotted into 96 well receivers from the same stocks for all studies. Spotted plates were stored at -80°C and thawed on the day of use. All tissues were fixed at 48 hours post infection (HPI). Candidate antiviral wells were normalized to control wells, included on each plate. The primary screen was performed with duplicate biological repeats. The final 11 candidate antivirals were tested in dose-response with triplicate biological replicates. All submerged and ALI plates were processed using a circular mask to remove false fluorescent signal from the edge of the wells. We also applied a size-based background removal on the ALI plates. All background removal steps were performed before the fluorescence of each well was measured and data was normalized. The research goal of 3D tissue studies was to develop a method to measure HSV-1 infection, candidate antiviral activity, and candidate antiviral toxicity in 3D bioprinted HSE.

All 2D monoculture experiments were controlled laboratory experiments using low passage primary keratinocytes and fibroblasts. HSV-1 aliquots were used a single time without refreezing. All candidate antivirals were serially diluted in PBS from 10mM stocks in DMSO and stored as aliquots at -20°C. All candidate antivirals were tested in technical duplicate, and each cell type for each donor was tested in biological triplicate for each candidate antiviral. A biological replicate is classified as an independent experiment using different assay preparations. Individual wells from all live cell imaging experiments were manually confirmed to be clear of autofluorescent debris, such as plastic particles, which would erroneously impact IncuCyte identification of fluorescent signal. Wells with autofluorescent debris were excluded from analysis. The primary research goal for 2D monoculture was to compare candidate antiviral potency and efficacy between cell types and against 3D bioprinted HSE.

### Commercial Cell Culture and Transduction

Neonatal human dermal fibroblasts (HDF_N_, Zen Bio DFN-F) were cultured in Dulbecco’s Modified Eagle Medium (DMEM, Gibco 11965) supplemented with 10% Fetal Bovine Serum (FBS) and 1% penicillin streptomycin. The fibroblasts were fluorescently labeled by transduction with lentiviral particles for fluorescent whole-cell labeling with tdTomato expression (Takara 0037VCT). A solution of polybrene and lentivirus at 20 multiplicities of infection (MOI) was added to the flask for 4 hours at 37°C, then virus solution was removed, and cells were expanded for 10 days, then frozen. Neonatal Normal Human Epithelial Keratinocytes (NHEK_N_, ScienCell 2100) were cultured in KBM Gold basal medium (Lonza, Cat. # 00192151) supplemented with the KGM Gold Keratinocyte Growth Medium BulletKit (Lonza, Cat. #00192152). All cell types were incubated at 37°C and 5% CO_2_.

### Donor Procured Primary Cell Isolation and Culture

This study was approved by the University of Washington Institutional Review Board. All primary cells were collected under IRB approval STUDY00004312. Written informed consent was collected from all participants.

Primary cell culture was completed using matched donor cells from 3mm punch biopsies collected as previously described (69). All cell culture medium contained 25mg streptomycin (Gibco 15140-122) and 25,000 units Penicillin (Gibco 15140-122). Keratinocytes were grown on collagen coated plates (Corning 35440) using DermaCult basal medium (Stem Cell Technologies 100-0501) with DermaCult keratinocyte expansion supplements (100–0502), 38µg Hydrocortisone (Stem Cell Technologies 07925), and 125µg Amphotericin B (Gibco 15290-019) added. The keratinocyte culture medium was supplemented with 10µM of the Rho Kinase inhibitor Y-27632 dihydrochloride (Stem Cell 72304) for the first six days after collection (70). Fibroblasts were grown on standard uncoated tissue culture vessels using Fibroblast Basal Medium 2 (PromoCell C-23220) with Fibroblast Growth Medium 2 supplement pack (PromoCell C-39320). All cells were maintained at 37°C in a humidified incubator under 5% CO_2_.

### Preparation of Dermal Base Hydrogel

Gelatin powder (Sigma G1890, final concentration 0.045mg/mL) was dissolved in a fibrinogen solution (Sigma F3879, final concentration 7.7mg/mL) in PBS, at 37°C. After complete dissolution of the gelatin, collagen I (Corning 354249, final concentration 4mg/mL) was added, and the solution was mixed thoroughly. 10X phosphate-buffered saline (PBS, Invitrogen AM9624) was added to buffer the gelatin/fibrinogen/collagen solution. The mixture was kept at 37°C and neutralized with 1N NaOH immediately before adding the fibroblasts.

Fibroblasts at 70% confluence were dissociated, harvested from the flask, counted with a Countess II FL hemocytometer, and then centrifuged at 650x g for 4 minutes. The pellet was resuspended in ∼10mL of the Dermal Base Hydrogel, at 2 million cells per mL, and mixed well by gentle pipetting. The solution was loaded into a 10mL syringe (Hamilton 81620) and chilled in the fridge for ∼5 minutes. Finally, a 0.42mm ID luer lock needle tip (Nordson 7018263) was placed on the syringe and the syringe was loaded onto the RegenHU 3DDiscovery bioprinter.

### 3D Printing the Dermal Bioink

A plunger-based bioprinting system was used to extrude the dermal base hydrogel solution in a cylindrical-crosshatched pattern made up of two 0.25mm thick layers with a diameter of 4.2mm directly on top of the membrane (8µm pore-size PET) of an HTS Transwell 96-well Permeable Support Plate (transwell plate, Corning 3374). The extrusion pattern was designed with the RegenHU BioCAD Software. To minimize evaporation effects, the hydrogel was not printed in any edge wells. After bioprinting was complete, the transwell was placed on a receiver plate filled with 250μL per well of serum-free DMEM supplemented with 5units/mL thrombin (Sigma T6884) at RT to allow the fibrinogen-fibrin conversion. After 1 hour, media in the receiver plates was changed to DMEM medium supplemented with 10% FBS, 1% penicillin streptomycin, and 0.025mg/mL aprotinin (Sigma A4529). The plates were incubated at 37°C and 5% CO_2_; media was replaced every 48 hours.

### Addition of Keratinocytes

7 days after bioprinting the dermal base, keratinocytes (NHEK_N_) at 70% confluence were dissociated, harvested from the flask, counted using a hemocytometer, pelleted at 650x g for 4 minutes, then resuspended in KGM medium at 0.4 million cells per mL. 50µL of the keratinocyte suspension (20,000 cells) was pipetted on top of the 3D bioprinted dermal base. Media in the receiver plates was changed to 250µL of epidermalization medium per well supplemented with 0.025mg/mL aprotinin (30, 31, 35). The plates were incubated at 37°C and 5% CO_2_; media was replaced every 48 hours.

### Air-Liquid Interface

7 days after adding the keratinocytes, the ALI model tissues were lifted to air liquid interface (ALI) using a custom 3mm high 3D printed adaptor made of SBS-format compliant material. The adaptor lifted the transwell insert, leaving only the bottom of the dermal base in contact with the media while the top of the tissues was exposed to air. The media in the receiver plate was replaced with 510µL of cornification medium supplemented with 0.025mg/mL aprotinin (30, 31, 35). Plates were incubated at 37°C and 5% CO_2_; media was replaced every 48 hours.

### HSV Infection in 3D Bioprinted HSE

All viral infection experiments utilized a GFP-expressing recombinant HSV-1 strain, K26 (37). To infect bioprinted HSE, HSV-1 stocks were thawed over ice for ∼1 hour then diluted in media for the submerged infection method or cornification media for the ALI infection method. The optimal MOI of HSV-1 for infection assays in the 3D bioprinted HSE was determined by diluting HSV-1 in cell growth media to 0.1, 1.0, and 10 MOI, then adding the virus onto the apical side of the submerged tissues one day after seeding keratinocytes. Z-stacks were captured at 4X magnification on an inverted microscope, and a maximum projection was used to quantitate GFP and tdTomato total fluorescence signal of each well at 24, 48, and 72 hours post infection (HPI). In the subsequent antiviral screening experiments, we used the HSV-1 at 0.1 MOI and fixed tissues at 48HPI.

We implemented two methods of infecting the 3D bioprinted HSE for antiviral screening. In the submerged method, the virus and the antivirals were added to the tissues one day after seeding the keratinocytes to the dermal base. The tissues were infected on the apical surface, but compounds were added to the media on the basal side of the tissues. In the ALI model, the virus and compounds were both added to the media on the basal side of the tissue four days after lifting to ALI to mimic viral reactivation.

### Compound Plate Preparation

For the screen, compounds were spotted into a 96-well receiver plates with the drug solutions at the appropriate stock concentrations in DMSO and frozen at -80°C until ready for use. The plate map used for the screen is shown in Figure 3A, with controls included on each plate including an inhibitor control (media -HSV-1), a negative control (HSV-1 + DMSO), and a known compound (HSV-1 + 0.2µM acyclovir).

### Histology

Tissues were fixed in 4% Paraformaldehyde for 24 hours then soaked in 30% sucrose for 24 hours at 4°C. The tissues were cut out from the transwell plate with a scalpel then embedded in Scigen Tissue Plus O.C.T. Compound (Fisher Scientific 23-730-571) at -20°C. The embedded tissues were sliced in 10µm sections with a CryoStar NX50 then mounted on positively charged slides. H&E staining was performed on the ThermoFisher Gemini stainer using the predefined H&E protocol. Using the Bond Fully Automated IHC Staining System, tissues were stained for primary antibodies keratin 10 and keratin 14 with Akoya Opal secondaries in 690 and 520 respectively. Slide images were taken in 10X with the Leica Aperio Versa 200.

### Imaging and Data Normalization of 3D Printed Tissues

Tissues were fixed in 4% Paraformaldehyde for 24 hours before washing with PBS. Fluorescent images were taken with a Molecular Devices ImageXpress High Content Reader using a 4X objective and excitation and emission wavelengths of 475-650nm for GFP and 544-570nm for tdTomato. Z-stacks of the images were taken at 25µm step size and a maximum Z-projections were made for each well using the MetaXpress Image Analysis.

The fluorescent signal for each tissue was measured using the MetaXpress Image Analysis software. To remove the fluorescent signal of the well edges, a circular mask of ∼4mm in diameter was created and only signal within the ‘circle mask’ was measured. For the ALI tissues an additional step was included to decrease background autofluorescence signal; a size-based inclusion threshold of 14.8µm or more was used to measure ‘cell-sized’ objects. (fig. S3) For both models, the GFP and tdTomato fluorescence intensity from the z-stack maximum projection was measured for each well. The intensity values were then normalized to the high and low signal controls using the equation:

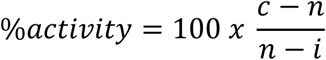

where *c* is the candidate antiviral fluorescence intensity value, *n* is the median signal from the negative controls (HSV-1 + DMSO), and *i* is the median signal of the inhibition control (media -HSV-1) for the GFP signal and 0 for the tdTomato signal.

### Analysis of 3D model applicability for high throughput screening

The reproducibility of results obtained using both the submerged and ALI 3D models was determined by Z-score, which was calculated using the following equation (38):

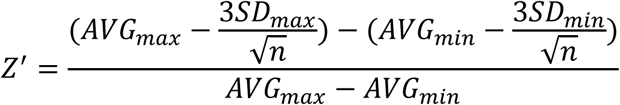

where *AVG_max_*=negative control (HSV-1 + DMSO) and *AVG_min_* =inhibitor control (media -HSV-1), *SD_max_* and *SD_min_* is the standard deviation of the negative and inhibitor control respectively, and *n* is the number of wells in each control group (38).

### 2D monoculture candidate antiviral testing

Primary keratinocytes were seeded in collagen coated 96 well plates (Revvity 6005810) and used at passage number six or less. Primary fibroblasts from the same keratinocyte donors were seeded in standard 96 well plates (Revvity 6005182) and used at passage number eight or less. 2D monocultures were infected with K26 HSV-1 using a MOI of 1.0 in serum free medium. Cells were infected at time zero and incubated on a rocker at 37°C and 5% CO_2_ for one hour prior to drug administration. Plates were imaged every two hours using a 10X objective on an IncuCyte S3/SX1. All images collected were analyzed using the IncuCyte software version 2021B (Sartorius). Infected wells treated with the 0.2% DMSO control were used to determine positive fluorescent signal while uninfected wells were used to determine background fluorescence for automated analysis. GFP expression is displayed as integrated intensity, defined as the total increase in fluorescent intensity as a function of the total area of detectable fluorescence.

Cytotoxicity was determined in the absence of HSV infection by treating cells with specified doses of candidate antivirals and using a combination of membrane permeable NucSpot Live 488 dye (Biotium 40081) and membrane impermeable NucSpot 594 dye (Biotium 41037). The cytotoxicity dose response curve was determined at 20 hours post treatment (HPT) for keratinocytes and 48 HPT for primary fibroblasts to align with IC_50_ timepoints.

### Plaque Reduction Assay

6-well plates (Corning 3516) were coated with 0.04mg/mL collagen I (Corning 354236) diluted in filter sterilized 0.1% acetic acid (VWR UN2789) for 1 hour, then seeded with primary keratinocytes. 95% confluent keratinocyte cultures were infected the next day using 40 plaque forming units of either HSV-1 K26 or HSV-2 186. Cells were infected at time zero and incubated on a rocker at 37°C and 5% CO_2_ for one hour prior to drug administration. DermaCult was prepared as described above then supplemented with 1% methylcellulose and candidate antivirals at the required concentrations. Excess virus was removed after one hour and replaced with overlay media containing candidate antivirals. Plaques were allowed to develop for 72 hours at 37°C and 5% CO_2_, then cells were fixed for 30 minutes in 4% paraformaldehyde (Boster Bio AR1068) and stained with crystal violet (J.T. Baker F906-03) before manual counting.

### IncuCyte IC_50_ Calculations

All IC_50_ values were determined by GraphPad Prism 9 using integrated fluorescent intensity of GFP measured by the IncuCyte software. IC_50_ values were calculated at 20, 36, or 48 HPI as specified. Uninfected wells served as the negative control and were used as the minimum signal (0%), while drug-free infected wells were used as the positive control and represent the maximum signal (100%). IC_50_ was calculated using a four parameter least squares nonlinear regression comparing the drug concentration to normalized response.

### Statistical Analysis

Percent activity and percent viability in the 3D HSE was determined by normalizing the candidate antivirals to the control wells. Statistical analysis was completed using either Welch’s two-tailed T-test or Wilcoxon two-sample test as described in figure legends.

Outliers in 2D monolayer studies were determined as values at least two logs higher than the mean. GFP fluorescence was normalized in 2D studies by subtracting fluorescence in mock infected wells and dividing by fluorescence in control untreated infected wells. Viability was determined in 2D studies by subtracting the total dead cells from the total cells to determine total live cells, then normalized by dividing the number of live cells in each experimental sample by the number of live cells in the 0.2% DMSO control at that timepoint. Statistical analysis for data sets involving multiple donors were analyzed by linear mixed models to account for donor to donor variability. Comparisons between submerged and ALI models were completed using linear models as a single cell source was used for all 3D tissues. Fold change calculations with 2D and 3D models used geometric means to account for the large differences between values.

## List of Supplementary Materials

Fig. S1-S15

Table S1-S7

Supplementary Spreadsheet S1-S5

## Acknowledgements

**Funding**: This work was supported by:

National Institutes of Health grant AI143773 (JZ)

National Institutes of Health TR003208 (JZ)

National Institutes of Health T32 AI07140 to (IRH)

NIH Intramural Research Program and Cure Acceleration Network Program to NCAT (MF)

## Author Contributions

S.T.E designed, performed, and analyzed the 3D bioprinted human skin equivalent studies and wrote and reviewed the manuscript. I.H. designed, performed, and analyzed the 2D studies and wrote and reviewed the manuscript. K.D. and P.D. developed the 3D bioprinted human skin equivalent protocol. S.F. designed and manufactured the custom 3D printed lifters. Z.I. prepared the compound plates for the 3D bioprinted human skin equivalent studies. M.S. performed data analysis on 3D bioprinted human skin equivalent studies. A.J. and W.O. designed, performed, and analyzed the 2D monoculture experiments. L.C. discussed the data and reviewed the manuscript. A.W. and C.J. oversaw clinic operation and donor sample collection and reviewed the manuscript. Y.F. performed the data and statistical analysis. M.F. conceived, designed, and supervised the 3D bioprinted human skin equivalent studies, wrote and reviewed the manuscript, and obtained funding support. J.Z. conceived, designed, supervised, and analyzed the study, wrote and reviewed the manuscript, and obtained funding support.

## Competing Interests

Authors declare that they have no competing interests

## Data and materials availability

All data are available in the main text or the supplementary materials.

## Notes

### Competing Interest Statement

The authors have declared no competing interest.

### Summary of Updates

Hello, we have updated the manuscript to highlight the importance of the limitations of the common HSV antiviral Acyclovir

