## Supplemental Figures for "Limitations of acyclovir and identification of potent HSV antivirals using 3D bioprinted human skin equivalents"

**SUPPLEMENTARY MATERIALS**

| Donor | Race and ethnicity | Age | Sex Assigned at Birth | HSV1 seropositivity | HSV2 seropositivity |
|---|---|---|---|---|---|
| 1 | White | 75 | Female | Positive | Positive |
| 2 | White | 62 | Female | Negative | Positive |
| 3 | White | 46 | Female | Positive | Positive |
| 4 | White, Hispanic or Latino | 29 | Male | Negative | Negative |
| 5 | White | 28 | Female | Negative | Negative |
| 6 | White, Hispanic or Latino | 30 | Female | Negative | Negative |

**Table S1. Donor demographic and HSV seropositivity information**

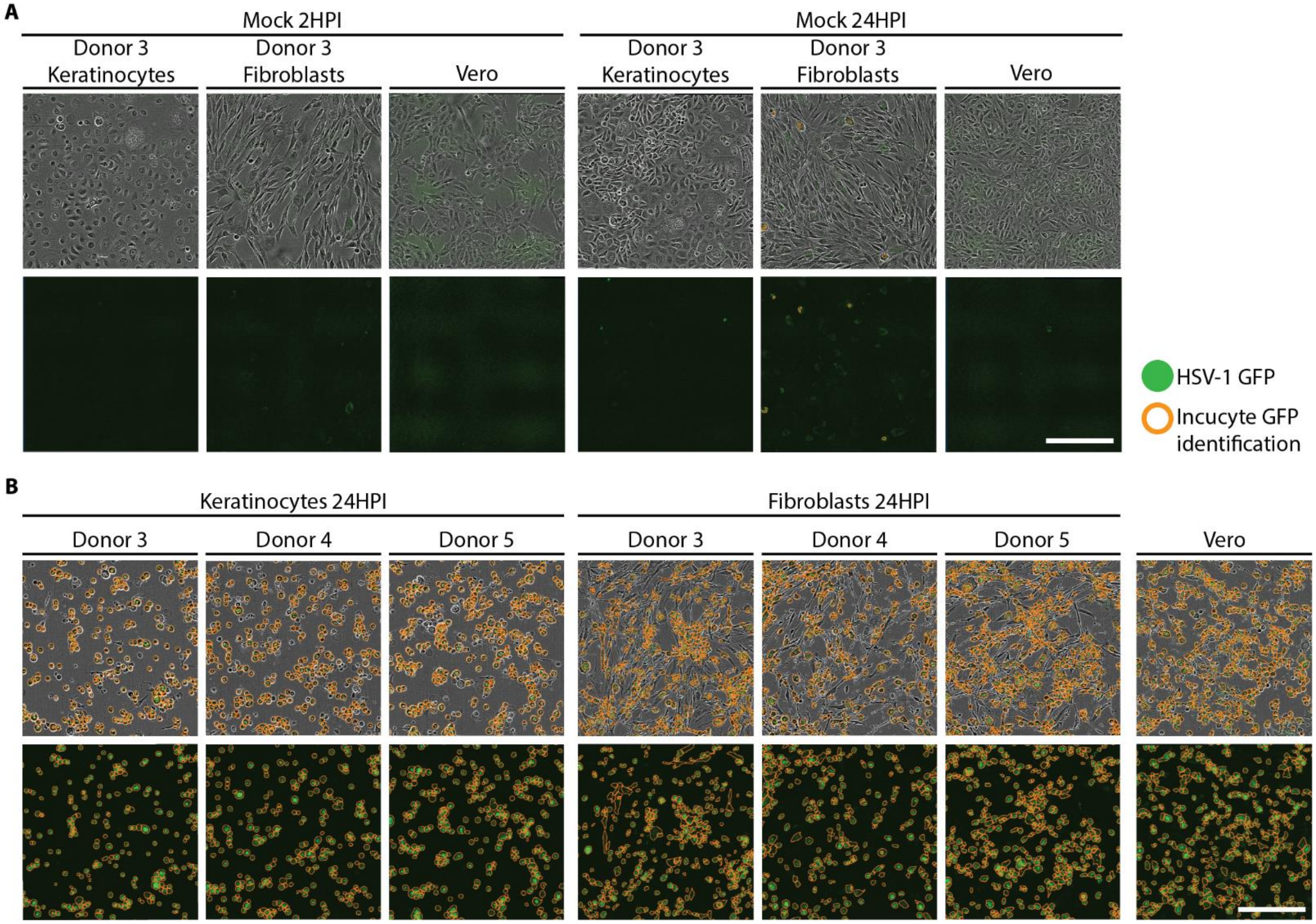


**Fig. S1. Validation of Incucyte automated identification of GFP fluorescence.**

**(A)** Representative images of mock infected cell cultures at 2 hours post infection (HPI) and 24HPI were used to establish background autofluorescence. **(B)** Representative images of infected Vero cells, keratinocytes, and fibroblasts at 24HPI were used to confirm that the Incucyte software successfully identified GFP positive cells (orange outlines). Scale bar is 250μM.

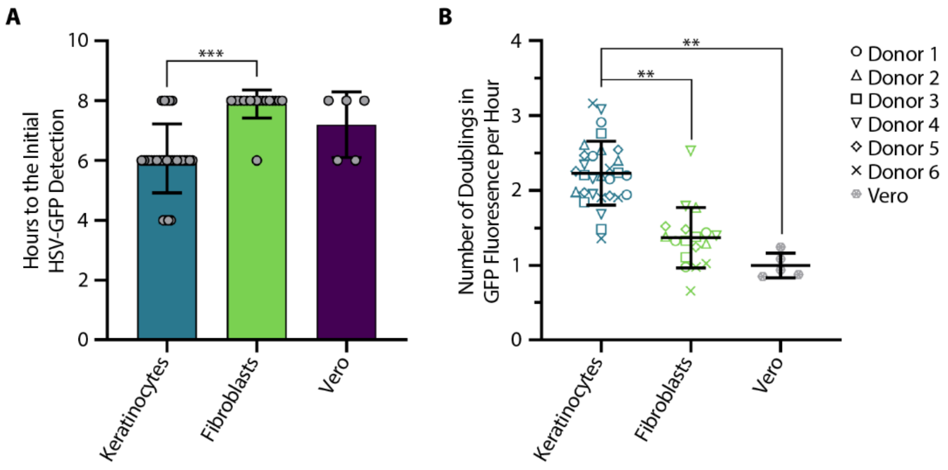


**Fig. S2. Virus Encoded GFP expression in primary keratinocytes and fibroblasts**

**(A)** Comparison of time until HSV-GFP was detected in 2D primary cell culture of specified cell types ( *** $P < 0.001$, Welch's T-test, $t = 7.87$, $df = 42.47$, keratinocytes ($N = 30$), fibroblasts ($N = 18$), and Vero cells ($N = 5$)). **(B)** Comparison of doubling time for virus-encoded GFP fluorescence in primary 2D monoculture ( ** $P < 0.01$). Keratinocytes vs Vero student's T-test, df= 5, keratinocytes ($N = 6$), Vero ($N = 1$). Keratinocytes vs fibroblasts Welch's T-test, $t = 6.02$, df 5.89, keratinocytes ($N = 6$), fibroblasts ($N = 6$). A value of 1 means GFP fluorescence doubled once per hour. Average for each cell type is plotted, individual biological replicates are represented by symbols, and error bars represent a single standard deviation.

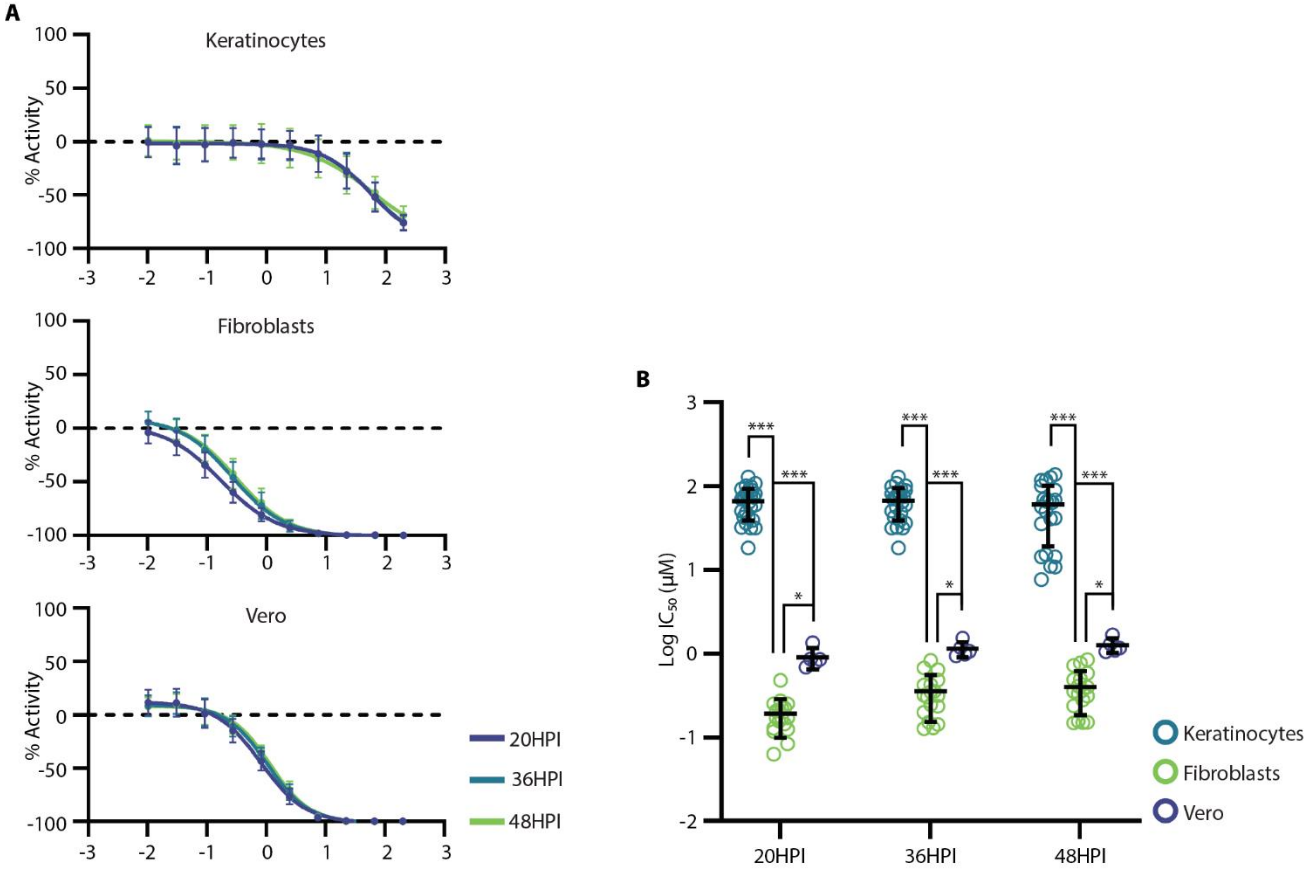


**Fig. S3. Acyclovir dose response and $IC_{50}$ in Vero cells, keratinocytes, and fibroblasts at 20, 36, and 48 hours post infection**

(**A**) Dose response curves for Vero cells, keratinocytes, and fibroblasts at each of the specified times. Cell type is specified in the y-axis of each plot. Average curves represent N = 30, 18, and 5 for keratinocytes, fibroblasts, and Vero cells respectively. Error bars are single standard deviations. **(B)** Calculated $IC_{50}$ values for acyclovir in each cell type at each of the specified times (* $P < 0.05$, *** $P < 0.001$, linear mixed model). Average for each cell type is plotted, individual biological replicates are represented by symbols and error bars represent a single standard deviation. Keratinocytes (N = 30), fibroblasts (N = 18), and Vero cells (N = 5) respectively

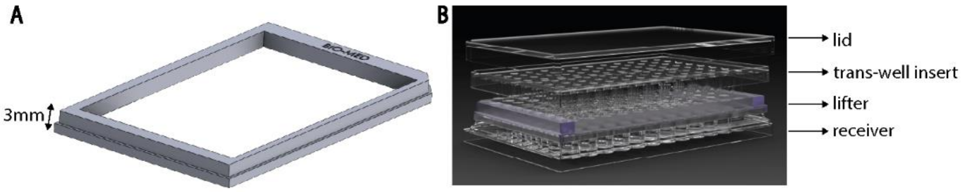


**Fig. S4. 3D printed lifter design for generating ALI cultures**

**(A)** Render of 3D printed lifter used to elevate 96-well transwell inserts to generate ALI cultures. **(B)** Depiction of lifter application within a standard 96-well transwell plate. Lifter is made of SBX-compliant material.

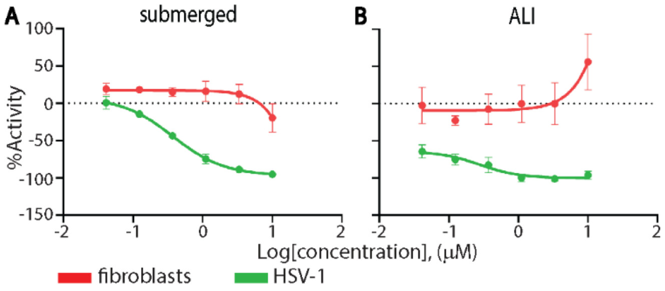


**Fig. S5. Dose response of acyclovir in submerged and ALI cultures**

**(A)** Acyclovir dose response (green) in the submerged model and in the **(B)** ALI model. Fibroblast transfected to express tdTomato signal (red) measures cytotoxicity. Average curves (N = 6) are plotted with error bars representing a single standard deviation.

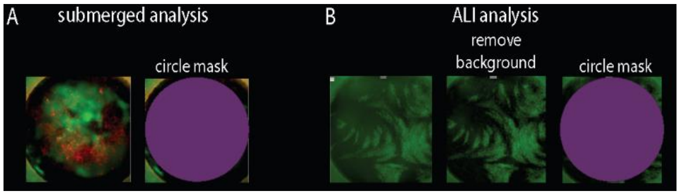


**Fig. S6. Background fluorescence signal mitigation in 3D bioprinted assay**

**(A)** In the submerged model, the false fluorescent signal of the plastic well edges is removed by using a circle mask. Only the signal within the circle mask is measured. **(B)** In the ALI model, first foggy background signal is removed using a size exclusion to eliminate sources of green fluorescence that were smaller than cells (filter 14.8 μm). After the background signal is removed, the same circle mask as in (A) is applied to mitigate the fluorescence of the well edges.

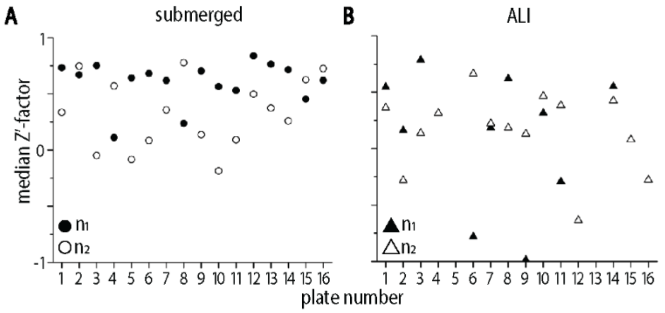


**Fig. S7. Median Z'- factor for submerged and ALI models**

**(A)** Median Z'-factor across each plate of 3D bioprinted tissues to measure statistical variance in the submerged model and the **(B)** ALI model. Each plate was completed in duplicate with n1 and n2 shown in closed and open symbols respectively.

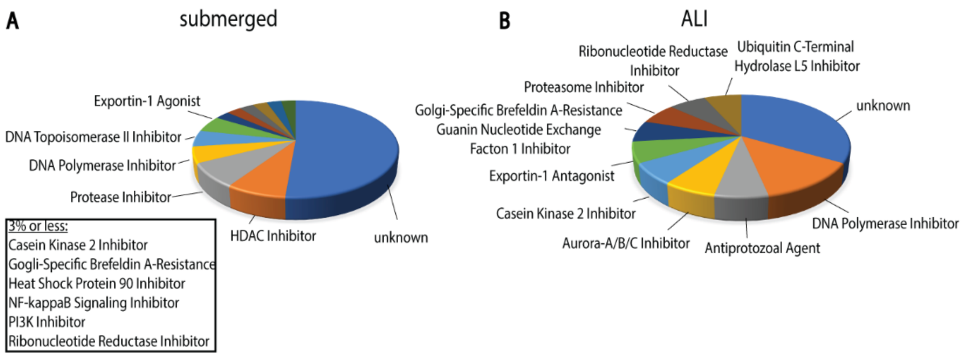


**Fig. S8. Secondary screen candidate antiviral mechanisms of action**

A proportion of known mechanisms of action for all 123 candidate antivirals in the secondary screen are displayed for submerged models **(A)** and ALI models **(B)**.

| Candidate antiviral | Literature |
|---|---|
| Acyclovir | Taylor and Gerriets, Acylcovir, 2023 |
| Adefovir dipivoxil | De Clercq, Clin Microbiol Rev, 2003 |
| Amenamevir | Chono et al, J Antimicrob Chemother, 2010 |
| Bardoxolone methyl | Wyler et al, Nat Commun, 2019 |
| Bortezomib (PS-341) | Schneider et al, mBio, 2019 |
| Cyanein | Ushio et al, Biomed Res, 2009 |
| Cycloheximide | Preston et al, J Gen Virol, 1998 |
| Emetine Dihydrochloride | Andersen et al, Viruses, 2019 |
| Epoxomicin | La Frazia et al, Antivir Ther, 2006 |
| Fiacitabine | Trousdale et al, Invest Ophthalmol Vis Sci, 1981 |
| Ganciclovir | Poole and James, Clin Ther, 2020 |
| Gemcitabine | Denisova et al, J Biol Chem, 2012 |
| GS7340 | Tan, Int J Womens Health, 2012 |
| Lanatoside C | Wu et al, Phytomedicine, 2024 |
| MG-132 | Ishimaru et al, Sci Rep, 2020 |
| Mitoxantrone | Huang et al, BMC Microbiol, 2019 |
| Niclosamide | Andersen et al, Viruses, 2019 |
| Penciclovir | Poole and James, Clin Ther, 2020 |
| Pritelivir (BAY 57-1293) | Betz et al, Antimicrob Agents Chemother, 2002 |
| SNX-2112 | Xiang et al, Bioorg Med Chem Lett, 2012 |
| Trifluorothymidine | Carmine et al, Drugs, 1982 |
| Valaciclovir | Spruance et al, Arch Intern Med, 1996 |
| Valganciclovir HCl | Poole and James, Clin Ther, 2020 |

**Table S2. Selected compounds that have published antiherpes activity.**

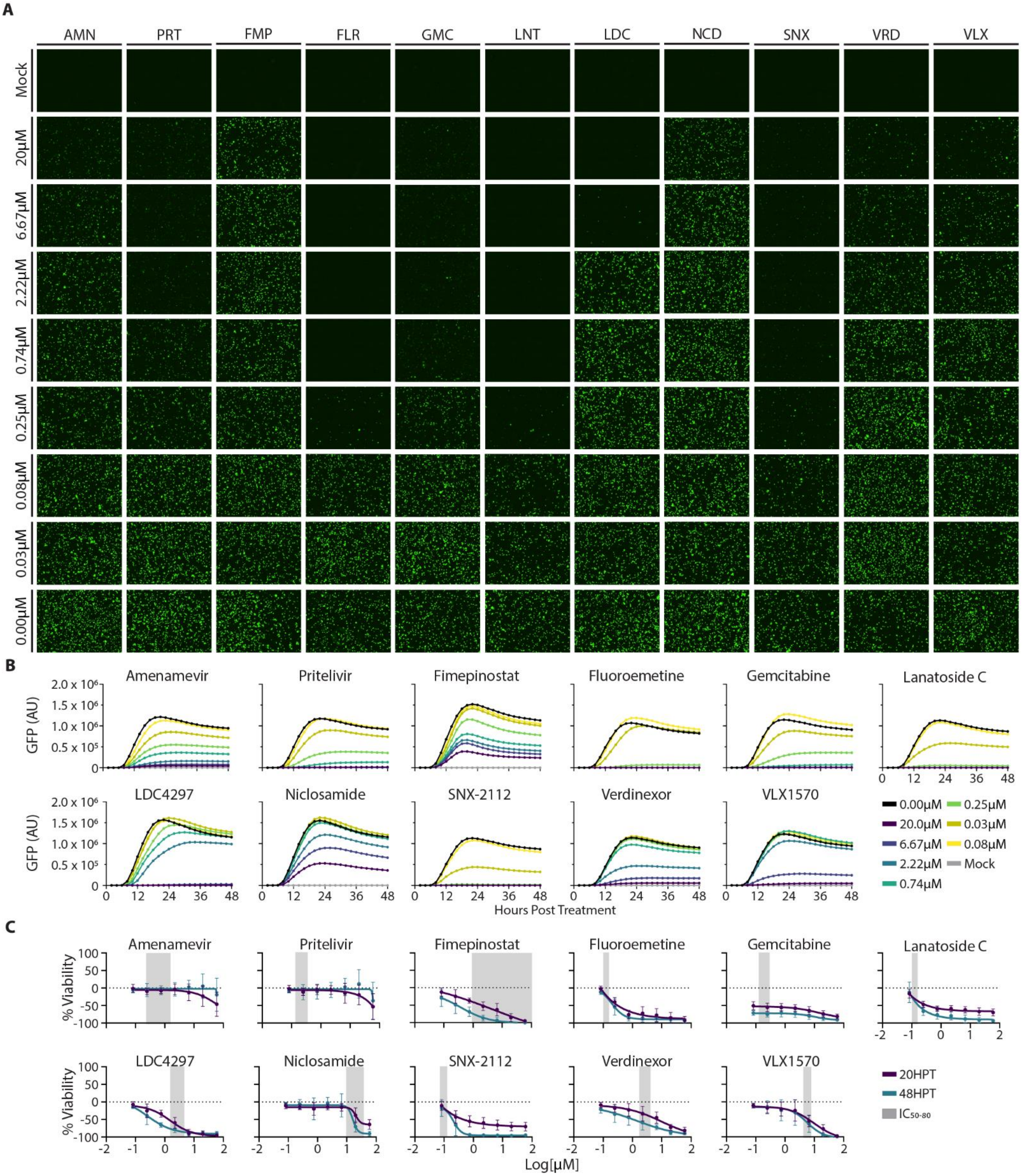


**Fig. S9. Dose response and cytotoxicity of candidate antivirals in donor-derived keratinocytes.**

**(A)** Live cell images of GFP expression during viral replication at 20HPI. Scale bar is 500μM. **(B)** Raw GFP integrated fluorescence in infected keratinocytes was calculated from live cell imaging collected every two hours. Each candidate antiviral was analyzed using 1:3 dilutions ranging from 20μM to 0.08μM. **(C)** Cytotoxicity of each candidate antiviral in uninfected cells at 20HPT (purple) and 48HPT (blue) compared to keratinocyte average $IC_{50}$ to $IC_{80}$ dose range (grey). All data represents average of three replicates for three donors (N = 9). Error bars represent a single standard deviation.

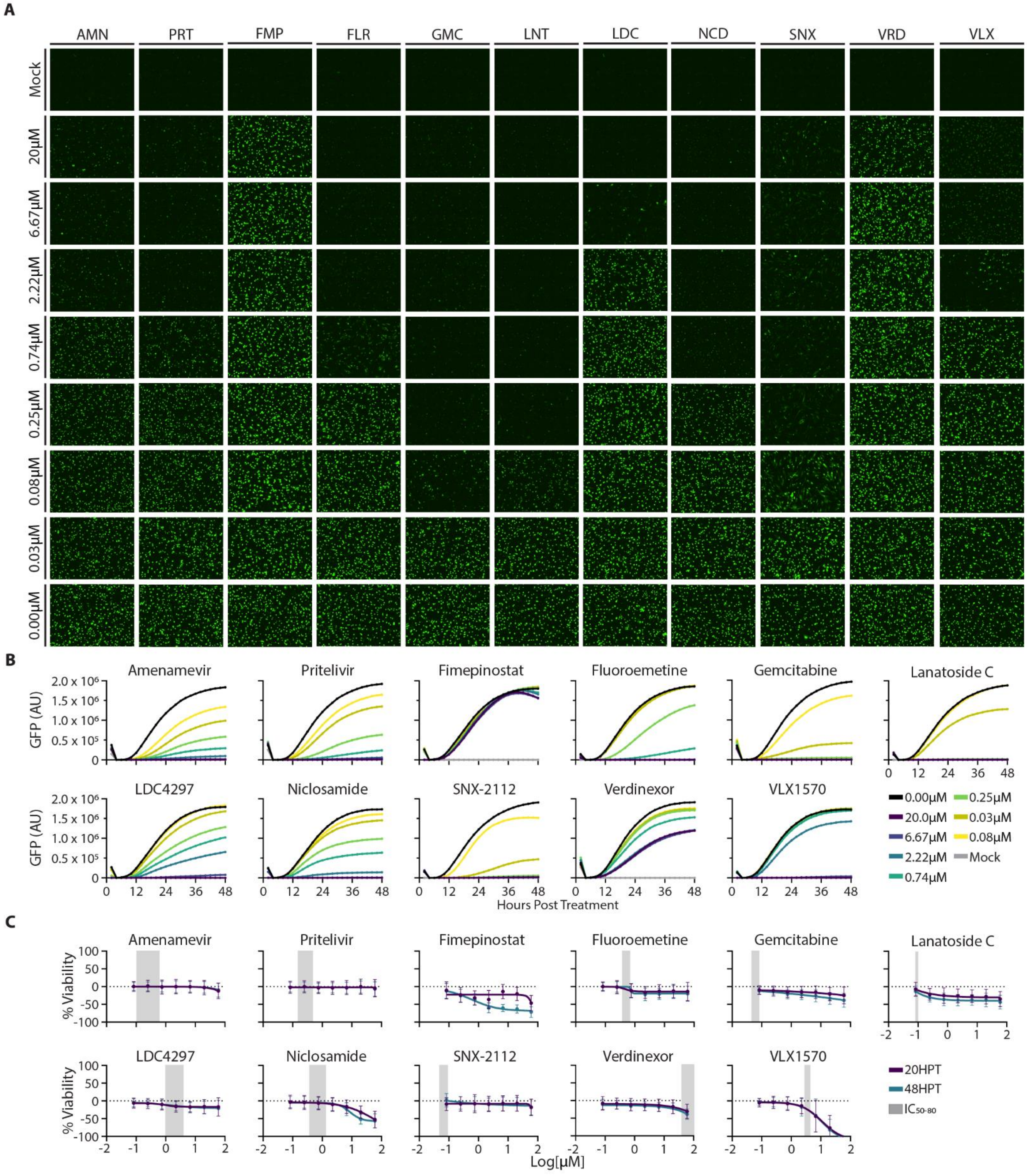


**Fig. S10. Dose response and cytotoxicity of candidate antivirals in donor-derived fibroblasts.**

**(A)** Live cell images of GFP expression during viral replication at 20HPI. Scale bar is 500μM. **(B)** Raw GFP integrated fluorescence in infected fibroblasts was calculated from live cell imaging collected every two hours. Each candidate antiviral was analyzed using 1:3 dilutions ranging from 20μM to 0.08μM. **(C)** Cytotoxicity of each candidate antiviral in uninfected cells at 20HPT (purple) and 48HPT (blue) compared to fibroblast average $IC_{50}$ to $IC_{80}$ dose range (grey). All data represents average of three replicates for three donors (N = 9). Error bars represent a single standard deviation.

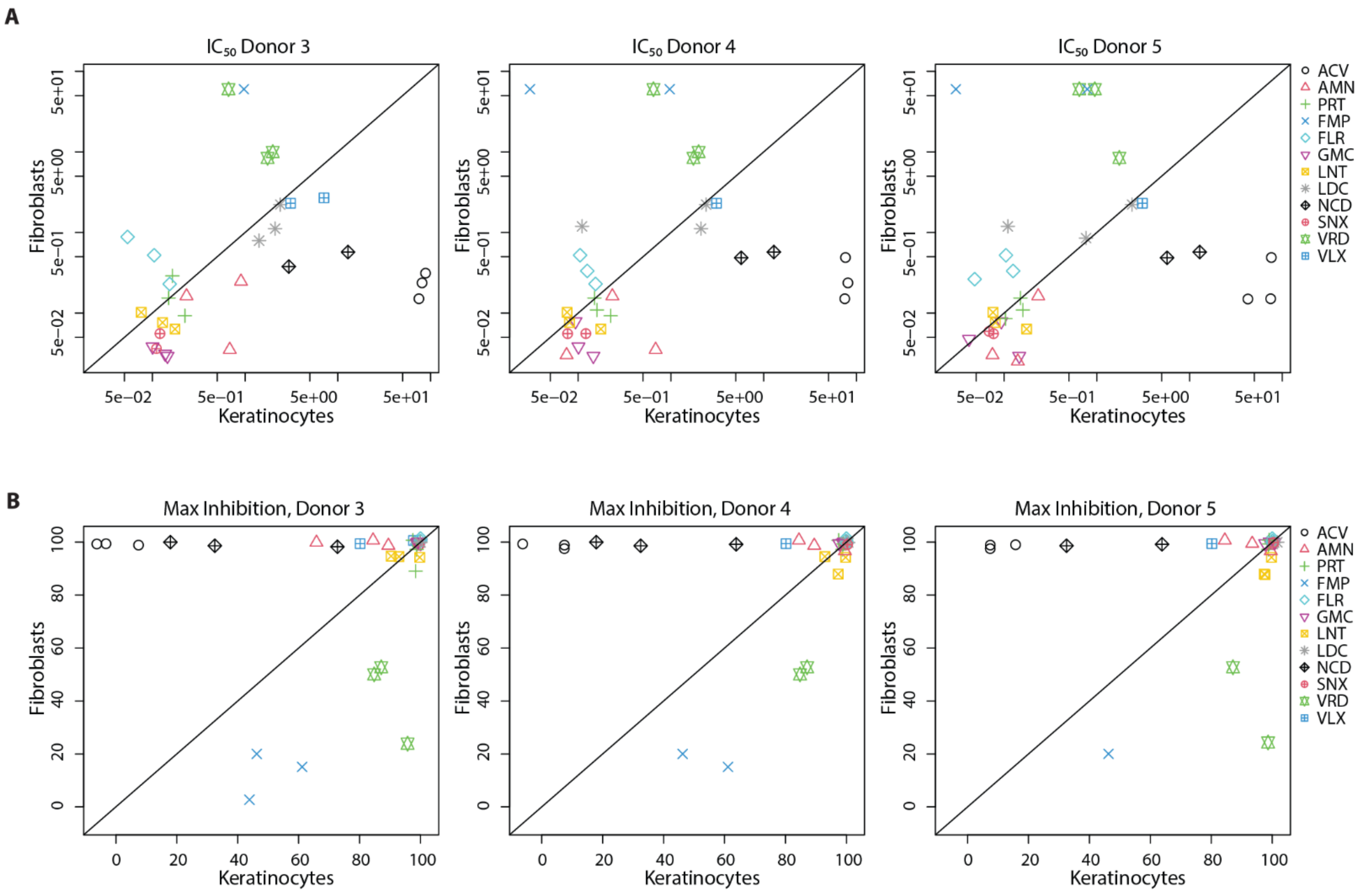


**Fig. S11. Pairwise comparisons of keratinocytes versus fibroblasts for each individual donor**

**(A)** Correlation graphs of Log [$IC_{50}$] values for keratinocytes (X axis) versus fibroblasts (y axis) for each donor. Symbols represent donor average (N = 3) **(B)** Correlation graphs of maximum inhibition at 10μM for each candidate antiviral in keratinocytes (X axis) versus fibroblasts (y axis) for each donor. Symbols represent donor cell type average (N = 3)

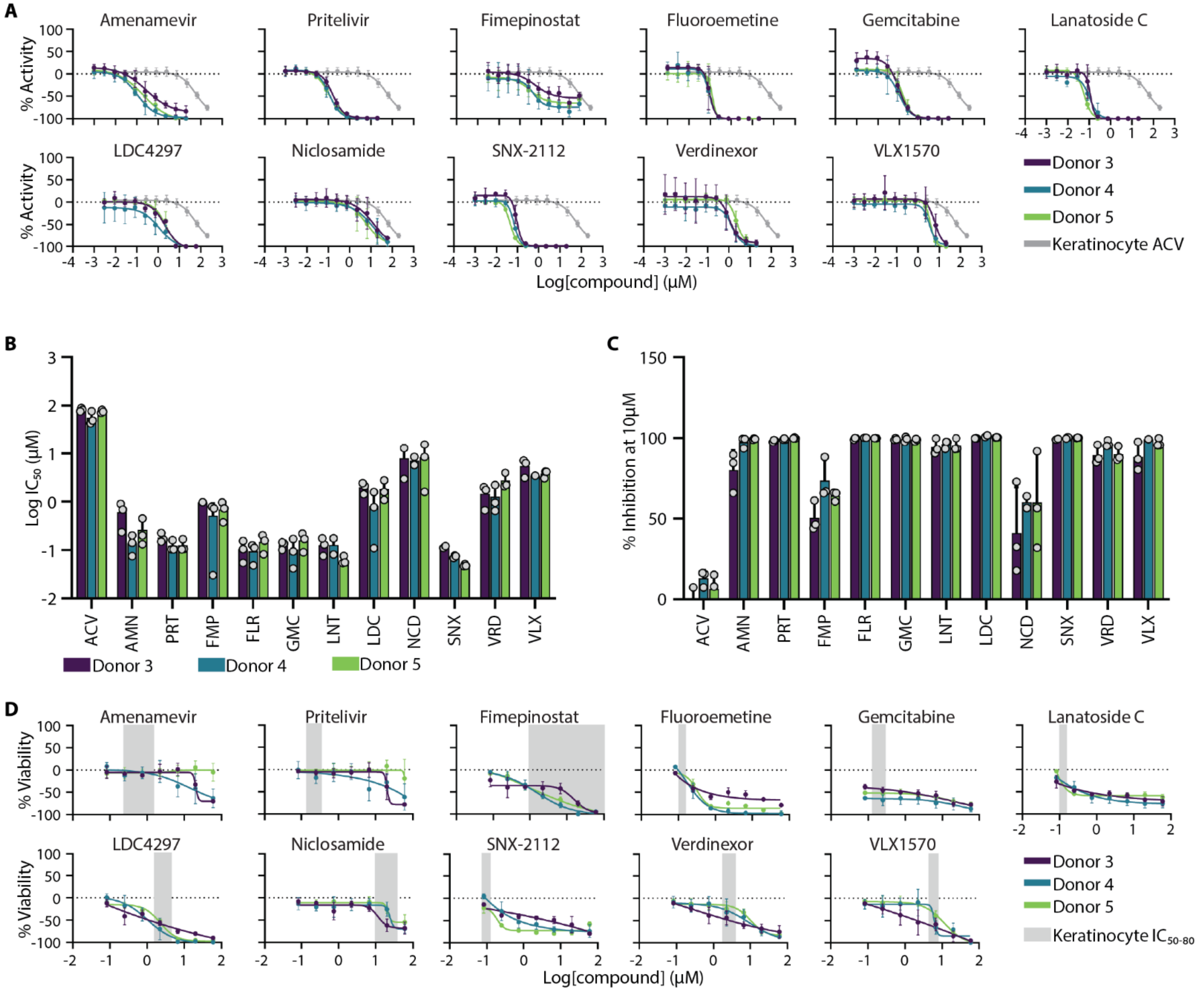


**Fig. S12. Candidate antiviral potency, efficacy, and cytotoxicity are comparable in keratinocytes between donors.**

**(A)** $IC_{50}$ dose response curves for all twelve top candidate antivirals compared to acyclovir (Donor 3 purple, Donor 4 blue, Donor 5 green, keratinocyte average ACV response grey). Curves repr **(B)** Absolute $IC_{50}$ values for each top candidate antiviral in keratinocytes is compared between Donor 3 (purple), Donor 4 (blue), and Donor 5 (green). **(C)** Total inhibition at 10µM for each top candidate antiviral is compared between Donor 3 (purple), Donor 4 (blue), and Donor 5 (green). **(D)** $CC_{50}$ dose response curves for all twelve candidate antivirals compared to their respective $IC_{50}$ to $IC_{80}$ dose range. Donor 3 is purple, Donor 4 is blue, Donor 5 is green, and average keratinocyte $IC_{50}$ to $IC_{80}$ dose range is grey. Abbreviations are acyclovir (ACV), amenamevir (AMN), pritelivir (PRT), fimepinostat (FMP), fluoroemetine (FLR), gemcitabine (GMC), lanatoside C (LNT), LDC4297 (LDC), niclosamide (NCD), SNX-2112 (SNX), verdinexor (VRD), and VLX1570 (VLX). All data represents average of three replicates (N = 3) for each donor. Error bars represent a single standard deviation.

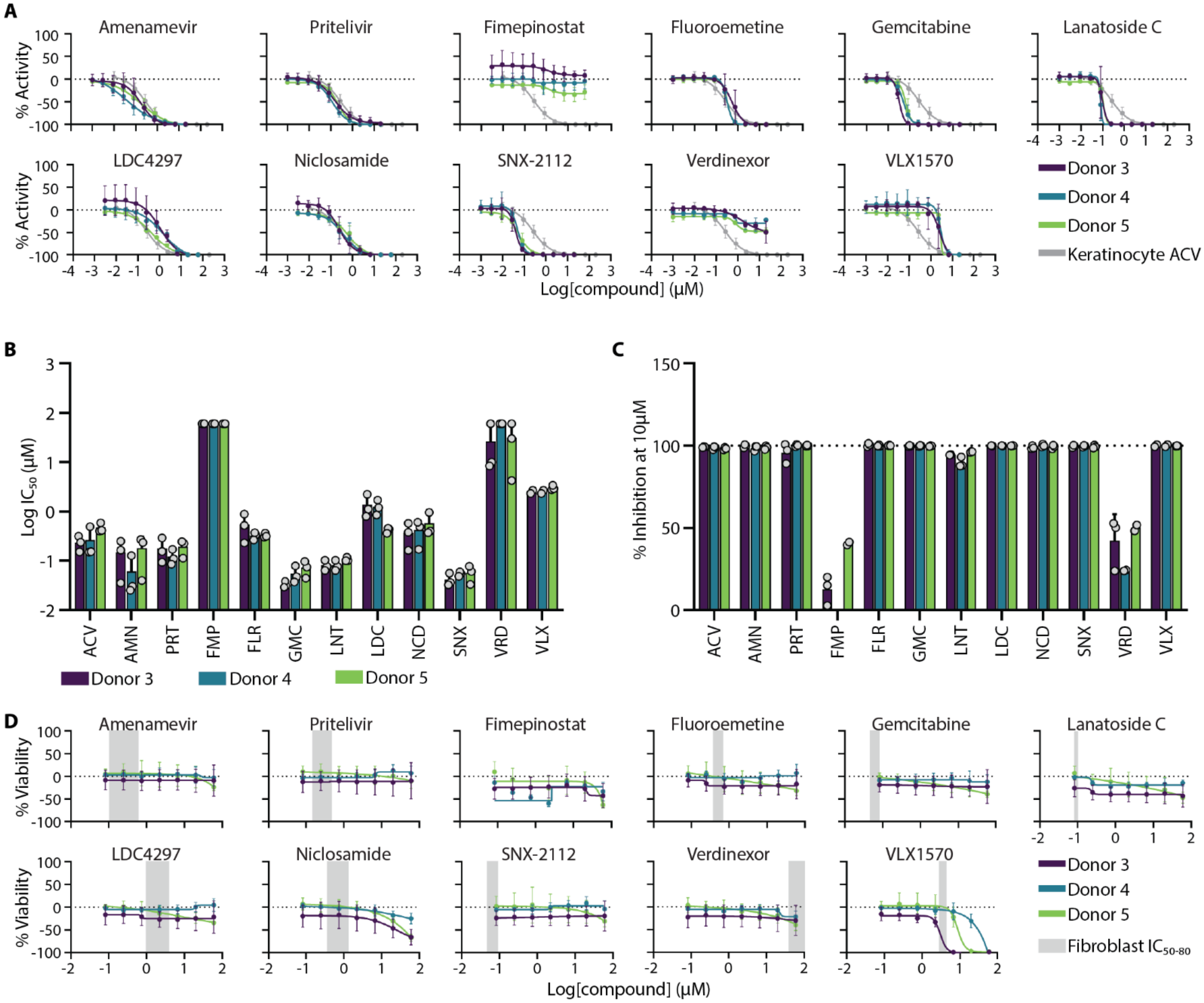


**Fig. S13. Candidate antiviral potency, efficacy, and cytotoxicity in individual donor-derived fibroblasts.**

**(A)** $IC_{50}$ dose response curves for all twelve top candidate antivirals compared to acyclovir (Donor 3 purple, Donor 4 blue, Donor 5 green, fibroblast average ACV response grey). **(B)** $IC_{50}$ values for each candidate antivirals in fibroblasts from three independent donors. Donor 3 (purple), Donor 4 (blue), and Donor 5 (green). **(C)** Total inhibition at 10μM for each top candidate antiviral is compared between Donor 3 (purple), Donor 4 (blue), and Donor 5 (green). **(D)** $CC_{50}$ dose response curves for all twelve candidate antivirals compared to their respective $IC_{50}$ to $IC_{80}$ dose range. Donor 3 is purple, Donor 4 is blue, Donor 5 is green, and the average fibroblast $IC_{50}$ to $IC_{80}$ dose range is grey. Abbreviations are acyclovir (ACV), amenamevir (AMN), pritelivir (PRT), fimepinostat (FMP), fluoroemetine (FLR), gemcitabine (GMC), lanatoside C (LNT), LDC4297 (LDC), niclosamide (NCD), SNX-2112 (SNX), verdinexor (VRD), and VLX1570 (VLX). All data represents average of three replicates (N = 3) for each donor. Error bars represent a single standard deviation.

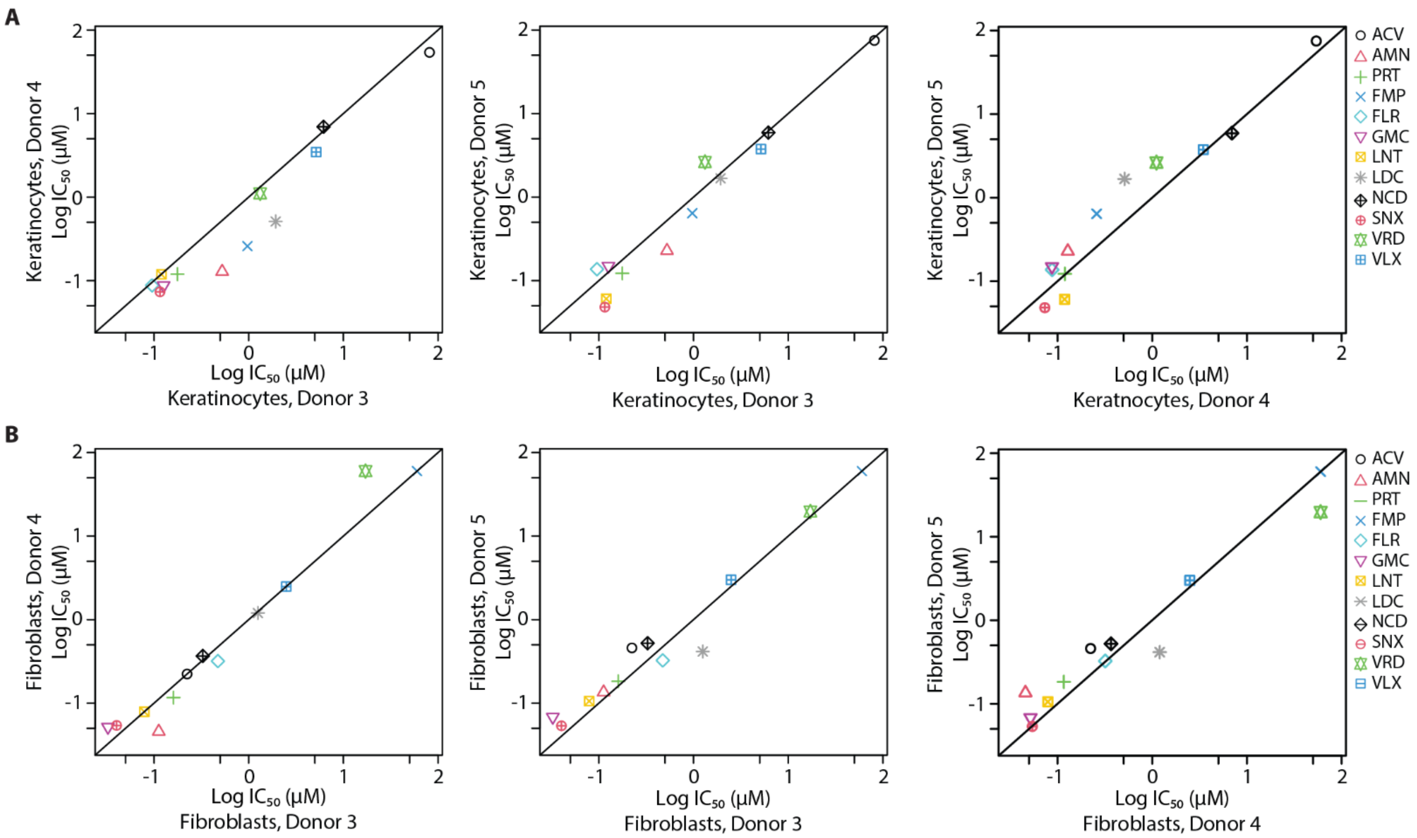


**Fig S14: Pairwise comparisons of candidate antiviral $IC_{50}$ values between donors**

Correlation graphs of absolute Log [$IC_{50}$] for each candidate antiviral in the specified donors for **(A)** keratinocytes and **(B)** fibroblasts. Symbols represent donor average (N = 3)

| | $IC_{50}$ | |
|---|---|---|
| | Keratinocytes | Fibroblasts |
| 3 versus 4 | < 0.001 | 0.768 |
| 3 versus 5 | 0.214 | 0.309 |
| 4 versus 5 | 0.024 | 0.471 |

**Table S3: P value results of pairwise comparisons**

Donor specific differences were calculated using a liner mixed model. Candidate antiviral potency is both cell types was determined as an absolute $IC_{50}$.

| | | Keratinocytes | Fibroblasts |
|---|---|---|---|
| Amenamevir (AMN) | $IC_{50}$ (µM) | 0.24 | 0.10 |
| | $CC_{50}$ (µM) | > 60.00 | > 60.00 |
| | Selectivity | > 250.00 | > 600.00 |
| Pritelivir (PRT) | $IC_{50}$ (µM) | 0.14 | 0.15 |
| | $CC_{50}$ (µM) | 55.50 | > 60.00 |
| | Selectivity | 396.43 | > 400.00 |
| Fimepinostat (FMP) | $IC_{50}$ (µM) | 0.95 | > 60.00 |
| | $CC_{50}$ (µM) | 1.97 | 1.49 |
| | Selectivity | 2.07 | < 0.02 |
| Fluoroemetine (FLR) | $IC_{50}$ (µM) | 0.10 | 0.38 |
| | $CC_{50}$ (µM) | 0.31 | > 60.00 |
| | Selectivity | 3.10 | > 157.89 |
| Gemcitabine (GMC) | $IC_{50}$ (µM) | 0.13 | 0.05 |
| | $CC_{50}$ (µM) | < 0.08 | > 60.00 |
| | Selectivity | < 0.62 | > 1200.00 |
| Lanatoside C (LNT) | $IC_{50}$ (µM) | 0.09 | 0.08 |
| | $CC_{50}$ (µM) | 0.33 | > 60.00 |
| | Selectivity | 3.67 | > 750.00 |
| LDC4297 (LDC) | $IC_{50}$ (µM) | 1.56 | 0.99 |
| | $CC_{50}$ (µM) | 1.36 | > 60.00 |
| | Selectivity | 0.87 | > 60.61 |
| Niclosamide (NCD) | $IC_{50}$ (µM) | 9.90 | 0.37 |
| | $CC_{50}$ (µM) | 24.47 | 19.48 |
| | Selectivity | 2.47 | 52.65 |
| SNX-2112 (SNX) | $IC_{50}$ (µM) | 0.08 | 0.05 |
| | $CC_{50}$ (µM) | 0.45 | > 60.00 |
| | Selectivity | 5.63 | > 1200.00 |
| Verdinexor (VRD) | $IC_{50}$ (µM) | 1.76 | > 60.00 |
| | $CC_{50}$ (µM) | 6.95 | > 60.00 |
| | Selectivity | 3.95 | 1.00 |
| VLX1570 (VLX) | $IC_{50}$ (µM) | 4.43 | 3.05 |
| | $CC_{50}$ (µM) | 6.35 | 8.34 |
| | Selectivity | 1.43 | 2.73 |

**Table S4: Comparison of IC50, CC50, and selectivity index for each top candidate antiviral in 2D monoculture**

$CC_{50}$ values of > 60.00µM indicate that 10µM doses did not achieve 50% loss of cellular viability. $CC_{50}$ values of < 0.08µM indicate that the lowest dose tested reduced the number of live cells by greater than 50%. $IC_{50}$ and $CC_{50}$ values represent the average of three replicates from each of the three donors (N = 9).

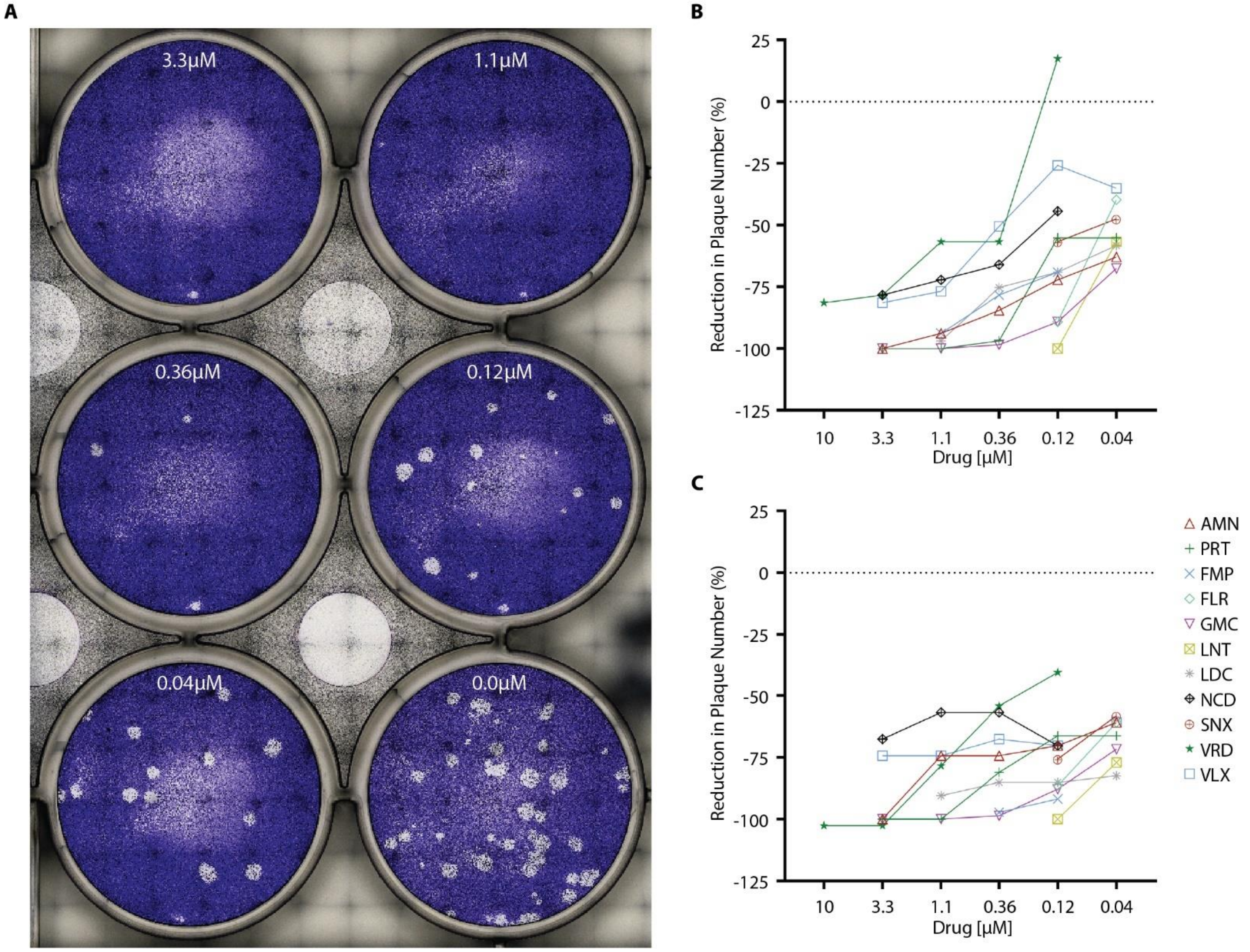


**Fig S15: Plaque reduction analysis of candidate antivirals against HSV-1 and HSV-2**

Primary keratinocytes were infected at 40 plaque-forming units per well of either HSV-1 K26 or HSV-2 186 with multiple doses of each candidate antiviral. (A) Example plaque reduction assay of pritelivir activity against HSV-2 186. (B&C) Percent of plaque reduction by candidate antivirals against HSV-1 K26 (B) or HSV-2 186 (C). Lines start at the highest dose without visible cytotoxic effects and end at lowest dose tested for each candidate antiviral. Data represents total plaques counted at each dose (N = 1).

| | | Submerged | ALI | Keratinocytes | Fibroblasts |
|---|---|---|---|---|---|
| Amenamevir (AMN) | $IC_{50}$ (µM) | 0.27 | 0.15 | 0.24 | 0.10 |
| | $CC_{50}$ (µM) | > 10.00 | > 10.00 | > 10.00 | > 10.00 |
| | Selectivity | > 37.04 | > 66.67 | > 41.67 | > 100.00 |
| Pritelivir (PRT) | $IC_{50}$ (µM) | 0.50 | 0.19 | 0.14 | 0.15 |
| | $CC_{50}$ (µM) | > 10.00 | > 10.00 | > 10.00 | > 10.00 |
| | Selectivity | > 20.00 | > 52.63 | > 71.43 | > 66.67 |
| Fimepinostat (FMP) | $IC_{50}$ (µM) | 1.45 | 0.04 | 0.95 | > 10.00 |
| | $CC_{50}$ (µM) | > 10.00 | > 10.00 | 1.97 | 1.49 |
| | Selectivity | > 6.90 | > 250.00 | 2.07 | < 0.15 |
| Fluoroemetine (FLR) | $IC_{50}$ (µM) | 0.15 | 0.22 | 0.10 | 0.38 |
| | $CC_{50}$ (µM) | > 10.00 | > 10.00 | 0.31 | > 10.00 |
| | Selectivity | > 66.67 | > 45.45 | 3.10 | > 26.32 |
| Gemcitabine (GMC) | $IC_{50}$ (µM) | 0.19 | 0.16 | 0.13 | 0.05 |
| | $CC_{50}$ (µM) | > 10.00 | > 10.00 | < 0.08 | > 10.00 |
| | Selectivity | > 52.63 | > 62.50 | < 0.62 | > 200.00 |
| Lanatoside C (LNT) | $IC_{50}$ (µM) | 0.09 | 0.08 | 0.09 | 0.08 |
| | $CC_{50}$ (µM) | 2.49 | > 10.00 | 0.33 | > 10.00 |
| | Selectivity | 27.67 | > 125.00 | 3.67 | > 125.00 |
| LDC4297 (LDC) | $IC_{50}$ (µM) | 0.67 | 0.11 | 1.56 | 0.99 |
| | $CC_{50}$ (µM) | > 10.00 | > 10.00 | 1.36 | > 10.00 |
| | Selectivity | > 14.93 | > 90.91 | 0.87 | > 10.10 |
| Niclosamide (NCD) | $IC_{50}$ (µM) | 0.36 | 0.11 | 9.90 | 0.37 |
| | $CC_{50}$ (µM) | > 10.00 | > 10.00 | > 10.00 | > 10.00 |
| | Selectivity | > 27.78 | > 90.91 | 1.01 | > 27.03 |
| SNX-2112 (SNX) | $IC_{50}$ (µM) | 0.05 | 0.04 | 0.08 | 0.05 |
| | $CC_{50}$ (µM) | 1.14 | > 10.00 | 0.45 | > 10.00 |
| | Selectivity | 22.80 | > 250.00 | 5.63 | > 200.00 |
| Verdinexor (VRD) | $IC_{50}$ (µM) | 0.35 | 0.15 | 1.76 | > 10.00 |
| | $CC_{50}$ (µM) | > 10.00 | > 10.00 | 6.95 | > 10.00 |
| | Selectivity | > 28.57 | > 66.67 | 3.95 | 1.00 |
| VLX1570 (VLX) | $IC_{50}$ (µM) | 6.30 | 0.15 | 4.43 | 3.05 |
| | $CC_{50}$ (µM) | 8.41 | > 10.00 | 6.35 | 8.34 |
| | Selectivity | 1.33 | > 66.67 | 1.43 | 2.73 |

**Table S5: Comparison of IC50, CC50, and selectivity index for each top candidate antiviral in 3D and 2D culture**

$CC_{50}$ values of > 10.00µM indicate that 10µM doses did not achieve 50% loss of cellular viability. $CC_{50}$ values of < 0.08µM indicate that the lowest dose tested reduced cellular viability by greater than 50%. $IC_{50}$ values of < 0.04µM indicate that the lowest does tested resulted in greater than 50% inhibition of virus encoded GFP expression. $IC_{50}$ and $CC_{50}$ values in 3D models represent the average of three replicates (N = 3) while in 2D models represent the average of three replicates from each of the three donors (N = 9).

| | Kera v. Fibro | Kera v. Sub | ALI v. Fibro | ALI v. Sub | 2D v. 3D |
|---|---|---|---|---|---|
| Acyclovir | 0.001 | 0.001 | 0.001 | 0.001 | 0.002 |
| Amenamevir | 0.032 | 1.000 | 1.000 | 0.602 | 1.000 |
| Pritelivir | 1.000 | 0.001 | 1.000 | 0.334 | 0.004 |
| Fimepinostat | 0.001 | 0.885 | 0.001 | 0.001 | 0.033 |
| Fluoroemetin | 0.001 | 1.000 | 0.235 | 0.412 | 1.000 |
| Gemcitabine | 0.001 | 0.909 | 0.034 | 1.000 | 0.033 |
| Lanatoside C | 1.000 | 1.000 | 1.000 | 1.000 | 1.000 |
| LDC4297 | 1.000 | 1.000 | 0.029 | 0.008 | 0.023 |
| Niclosamide | 0.001 | 0.001 | 0.001 | 0.167 | 0.033 |
| SNX-2112 | 0.104 | 1.000 | 1.000 | 1.000 | 0.440 |
| Verdinexor | 0.001 | 0.044 | 0.001 | 1.000 | 0.001 |
| VLX1570 | 0.005 | 0.832 | 0.001 | 0.004 | 0.122 |

**Table S6: P values comparing the IC50 of each model tested**

Candidate antiviral potency was compared between models using a linear mixed model. To compare 2D versus 3D, all potency values for each candidate antiviral in keratinocytes and fibroblasts were pooled, then compared to all potency values pooled for submerged and ALI models of the same candidate antiviral.

| | | Neonatal IC50 | Adult IC50 |
|---|---|---|---|
| Submerged | AMN | 0.27±0.04 | 0.38 |
| | PRT | 0.50±0.23 | 0.24 |
| | FMP | 1.48±0.99 | 0.37 |
| | FLR | 0.15±0.02 | 0.13 |
| | GMC | 0.19±0.08 | 0.16 |
| | LNT | 0.09±0.01 | 0.17 |
| | LDC | 0.68±0.36 | 0.70 |
| | NCD | 0.40±0.49 | 0.18 |
| | SNX | 0.06±0.07 | 0.09 |
| | VRD | 0.48±0.98 | 0.52 |
| | VLX | 6.67±7.17 | >10.00 |
| ALI | AMN | 0.16±0.22 | 0.10 |
| | PRT | 0.21±0.24 | 0.25 |
| | FMP | < 0.04±0.00 | 0.24 |
| | FLR | 0.22±0.14 | 0.28 |
| | GMC | 0.17±0.13 | 0.11 |
| | LNT | 0.09±0.04 | 0.14 |
| | LDC | 0.11±0.07 | 0.09 |
| | NCD | 0.11±0.04 | 0.25 |
| | SNX | < 0.04±0.00 | < 0.04 |
| | VRD | 0.17±0.24 | 0.26 |
| | VLX | 0.16±0.19 | 0.24 |

**Table S7: Comparison of IC50 values for neonatal versus adult-derived HSEs**

95% confidence intervals were calculated for each candidate antiviral IC50 using three biological replicates in neonatal keratinocyte derived HSEs (N = 3). Candidate antiviral IC50s in adult keratinocyte derived HSEs was determined from a single biological replicate (N = 1).
